# Lineage-wide evolution of 3D genome organisation and centromeres in brown algae

**DOI:** 10.64898/2025.12.07.692804

**Authors:** Pengfei Liu, Rory J. Craig, Elena Avdievich, Fabian B. Haas, Chang Liu, Susana M Coelho

## Abstract

Although 3D genome architecture has been described for an increasing number of plant and algal species, comparative analyses across closely related lineages remain scarce. Consequently, fundamental questions persist about how chromatin organization is maintained or reshaped over deep evolutionary time, and how such changes relate to life-history traits, genome size, and linear genome features. Here, we present a comprehensive analysis of 3D chromatin architecture across six brown algae species and one outgroup, spanning the phylogenetic breadth and biological complexity of this key photosynthetic lineage. We show that compact genomes lack chromatin domains whereas larger, transposable element-rich genomes of morphologically complex taxa tend to exhibit structured organization including TAD-like domains. We investigate chromatin folding patterns and gene expression over evolutionary time and uncover 3D chromatin features associated with transitions in sexual systems. Moreover, we reconstruct the ancestral brown algal karyotype, revealing deeply conserved macrosynteny and providing a new framework for interpreting chromosome-scale genome dynamics. Finally, we uncover an ancient and highly conserved association between centromeres and chromodomain-encoding retrotransposons, revealing a remarkable example of convergence in centromere–transposon co-evolution between brown algae and angiosperms, and one of the most stable examples of centromere-linked transposable elements known in eukaryotes. Together, our findings elucidate the evolutionary history of 3D chromatin and linear genome architectures across an entire eukaryotic lineage and highlight extreme centromere stability in brown algae, providing a powerful point of comparison with land plants and deepening our understanding of genome evolution in independent multicellular lineages.

**One sentence summary:** We present a lineage-wide evolutionary analysis of 3D genome architecture and organization across brown algae, revealing conserved chromosomal features, lineage-specific rearrangements, and long-term co-evolution of centromeres and retrotransposons that together illuminate how nuclear architecture evolves over hundreds of millions of years.

## Introduction

Eukaryotic genomes are organized within the 3D nuclear space, and this spatial architecture is fundamental to chromatin folding and genome regulation^1,2^. Advances in chromatin conformation capture technologies, particularly Hi-C, have transformed our understanding of higher-order genome organization across cell types and species ^3–8^ revealing conserved and divergent principles of chromosome folding. Current models of nuclear architecture recognize that chromosomes in the cell nucleus are organized as chromosome territories (CTs). At finer scales, genomes are partitioned into A/B compartments, topologically associating domains (TADs), and specific promoter– enhancer loops^1,9^ providing a multilayered regulatory framework that links 3D structure to gene activity.

The 3D nuclear architecture has been characterized in model plants^10–13^, animals^3,9,14–17^, and yeast^4,5,18^, yet our understanding of chromatin organization across broader eukaryotic lineages remains limited^19^. Moreover, while an increasing number of studies describe the 3D genome of individual organisms^1,20^, comparative studies examining the evolutionary dynamics of 3D genome architecture across related species within a lineage are notably scarce, leaving fundamental questions about the conservation and divergence of higher-order chromatin structure largely unresolved. Specifically, the degree to which 3D genome organization is maintained across species boundaries, and the potential correlations between architectural modifications and key life history traits, genome size variation, and other structural genomic changes, remain poorly understood.

In this context, the brown algae (Phaeophyceae) offer an interesting study system. They are the third most complex multicellular eukaryotic lineage^21^ having arisen independently from animal and plant lineages^22^, although they are photosynthetic organisms that have been extensively used for comparative studies with plant lineages^23,24^. Brown algae as a group present a range of levels of morphological complexity, sexual system and sexual dimorphism, and lengths of haploid and diploid phase of development^28^. Like land plants, brown algae undergo a haploid–diploid alternation of generations, and many species possess sexual systems reminiscent of bryophytes, with UV sex chromosomes and separate sexes^23,24,29,30^, although some have transitioned to hermaphroditism (monoicy). These life-history traits are expected to strongly influence genome evolution and 3D genome architecture, yet direct empirical evidence linking them to chromatin organization remains scarce.

Despite the recent sequencing of 65 genomes of brown algae^26^, the 3D architecture of the brown algal nucleus is largely unknwon, except for a recent study using the model brown alga *Ectocarpus* sp.7^31^. *Ectocarpus* is a simple, filamentous brown algae with a compact 200 Mbp^32,33^ genome. Hi-C analyses showed that *Ectocarpus* genome can be partitioned into loose or compact structural domains that bear some similarities to those of the mammalian A/B compartments. *Ectocarpus* interphase chromatin exhibits a non-Rabl 3D chromatin conformation, with strong contacts among telomeres and among centromeres, which feature centromere-specific LTR retrotransposons{Citation}. However, *Ectocarpus* chromosomes do not contain large local interactive domains (i.e., TADs), which are a predominant feature of animal genomes^34^ and large genomes of plants^35,36^. However, Ectocarpus lacks large local interactive domains (TADs), which are prominent in animals and large plant genomes, leaving open the question of whether its 3D genome structure is representative of the lineage.

Here, we present a comparative analysis of 3D chromatin architecture across six brown algal species plus one outgroup, spanning a range of sexual systems, genome sizes, and morphological complexity. We find that compact genomes lack chromatin domains, whereas larger, transposable element–rich genomes exhibit structured organization, including TAD-like domains, linking genome complexity with 3D architecture. Sex chromosome 3D structure is maintained during transitions to hermaphroditism, reflecting linear conservation of male-specific features and demonstrating long-term stability of chromosome-level architecture. Using new chromosome-scale assemblies, we reconstruct ancestral karyotypes, revealing deeply conserved macrosynteny alongside lineage-specific rearrangements. Finally, we characterize brown algal centromeres, uncovering their long-term co-evolution with centrophilic retrotransposons and remarkable conservation of centromeric regions over ∼240 million years. These findings position brown algae as a powerful comparative system for understanding the evolution of 3D genome organization and centromere biology, providing insights highly relevant to plant genomics and evolution.

## Results

### Hi-C-guided chromosome level assemblies reveal conserved macrosynteny

To assess the evolution of chromosome structure across a diverse set of brown algae, we utilized Hi-C-guided, chromosome-level assemblies for six brown algal genomes and one sister outgroup (**Table S1, S2**). These brown algal species represent four major orders, encompassing the phylogenetic history of the lineage^21,37^ as well as its diversity in terms of sexual systems, sexual dimorphism, morphological complexity, and types of life cycle^28^ (**Fig. 1A-B**). We built on the recently published chromosome-level assembly of *Ectocarpus* sp. 7^31^, and improved the assemblies of five recently published algal genomes^25,26^ to chromosomal-level using Hi-C (see methods): *Chordaria linearis, Undaria pinnatifida, Desmarestia herbacea, Desmarestia dudresnayi,* and *Dictyota dichotoma*. Furthermore, we produced *de novo* chromosomal-level assemblies using Nanopore and Hi-C technologies for *U. pinnatifida* and *D. dichotoma* female strains (**Fig. 1B, Fig. S1, Table S1, S2**, see methods), enabling comparison between male and female genomes for these dioicous species. The new assemblies show high DNA pairwise alignment identity compared to previous assemblies (**Fig. S2**) and yield genome sizes from 185.8 Mb (*Ectocarpus* sp.7^31^) to 848.5 Mb (*D. dichotoma*), with N50 values from 6.91 Mb to 28.1 Mb and BUSCO scores ranging from 87.1% to 89.0% completeness using the eukaryota_odb10 database^38^ (**Fig. S1, Table S2**). The differences in genome size are mainly explained by expansions of transposable elements (TEs) in species with larger genomes (**Fig. 1B, S3**).

**Fig. 1.**
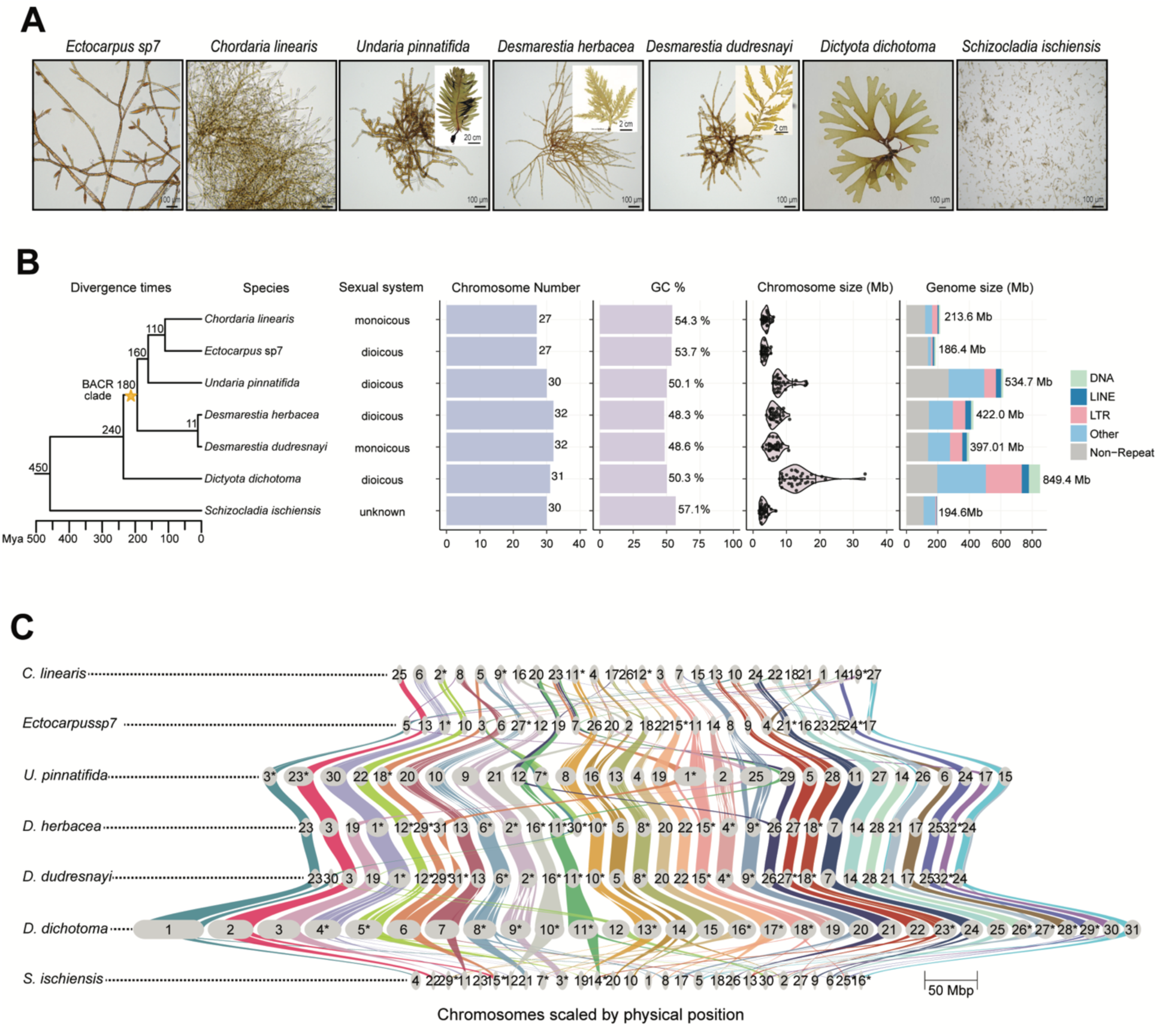
High quality genome assemblies for representative brown algal species and outgroup. **(A)** Species used in this study, covering the phylogenetic and morphological diversity of brown algae. Note that the images shown represent the gametophyte (haploid) generation whereas the inserts represent the morphologically complex sporophyte generation (diploid). **(B)** Phylogenetic position of the studied species and genome statistics. The phylogeny was built in TreeViewerV2.20^39^; the nodes are labelled with the approximate divergence time from^26^, the BACR (brown algal crown radiation) clade is marked with golden brown star. Repeats were grouped into major classes (DNA, LTR, LINE)^40^. All remaining categories, including unclassified repeats, simple repeats, satellites, and others, were combined into ‘other’. (**C**) Ribbon plot of syntenic orthologous genes conserved among brown algae and outgroup. The colored vertical links connect orthologous genes (identified using GENESPACE^41^) to the numbered chromosomes among the species. The order and color the chromosomes are rephased by *D. dichotoma*, and inverted chromosomes are label with an asterisk.

We also achieved a near-contiguous chromosome-level assembly for the sister group species *Schizocladia ischiensis* (**Table S2**) which has, as previously suggested, a highly rearranged genome relative to brown algae^25^. For synteny analyses, we focused on orthologs that were present on chromosomes (and excluded unlinked scaffolds, see methods) (**Fig. 1C, Fig. S1, Table S2**). Brown algae have 27-32 chromosomes and largely conserved macrosynteny and gene collinearity even after app. 240 million years of independent evolution (**Fig. 1B-C**), often differing by a few discrete events superimposed on a background rate of small-scale gene transfers between chromosomes. The evolutionary history of brown algal linear genomes is analyzed in greater detail below.

### 3D genome architecture of brown algae genomes

To uncover how 3D genome organization has evolved in brown algae, we utilized the new Hi-C datasets for the five focal species and one sister outgroup, and compared them with the published Hi-C map of the model alga *Ectocarpus*^31^ (**Table S1**). In a first step, we focused on the Hi-C maps from male individuals to perform the comparison across species. We then included male and female samples from dioicous species to examine potential sex differences in the 3D genome, together with derived monoicous (hermaphrodite) species to study the consequences of transitions to monoicy (and therefore loss of sex chromosomes) (**Table S2-S4**).

Biological replicates of Hi-C experiments were generated for each species and sex (**Fig. S4**). In total, we obtained between 263.5 and 892.5 million Hi-C interaction read pairs, depending on genome size and library efficiency (**Table S4**). Quality assessments showed strong reproducibility between biological replicates, with consistent cis-interaction frequency patterns (**Fig. S5A**) and Stratum-Adjusted Correlation Coefficient (SCC) scores^42^ (**Fig. S5B**). Distance-dependent contact frequencies (P(s)) and slope of P(s)^43^ reveal similar decay patterns for both replicates in all cases (**Fig. S5C, S5D**). Given the high reproducibility observed among biological replicates, the datasets were merged to maximize sequencing depth, thereby enabling downstream comparative analyses at higher resolution (**Fig. S6)**.

We examined the global chromatin compartmentalization across species using Hi-C contact maps. In the 3D nuclear space, genomes are typically organized into A (active) and B (inactive) chromatin compartments^3^. We derived A/B compartment profiles for each species by performing eigen- vector decomposition (**Fig. 2A, Fig. S7**) which showed that all species display compartmentalized chromosomes. Previous studies have shown that compartmentalization strength can vary across organisms and resolution scales^44^. Given the range of genome sizes among the species analyzed, we asked whether genome size influences compartment strength in brown algae. Compartment strength analysis using saddle plots (see Methods; **Fig. 2B**) revealed that the morphologically complex alga, *U. pinnatifida* (BB: 2.21, AA: 1.24; genome size: ∼534 Mb) and *D. herbacea* (BB: 2.10, AA: 1.18; genome size: ∼422 Mb), exhibited the strongest compartmentalization. In contrast, *S. ischiensis*, the morphologically simple outgroup species, showed the weakest compartmentalization (BB: 1.45, AA: 1.07; genome size: ∼195 Mb). *Ectocarpus* sp. 7 (BB: 1.62, AA: 1.56; genome size: ∼186 Mb) and *D. dichotoma* (BB: 1.47, AA: 1.01; genome size: ∼848 Mb) displayed comparable, intermediate compartment strengths despite their markedly different genome sizes. This contrast indicates that compartmentalization strength does not scale with genome size (**Fig. 2C, D**). Together, these results suggest that compartment strength is primarily shaped by species-specific genome organization and may be linked to increased morphological complexity rather than genome size alone.

**Fig. 2.**
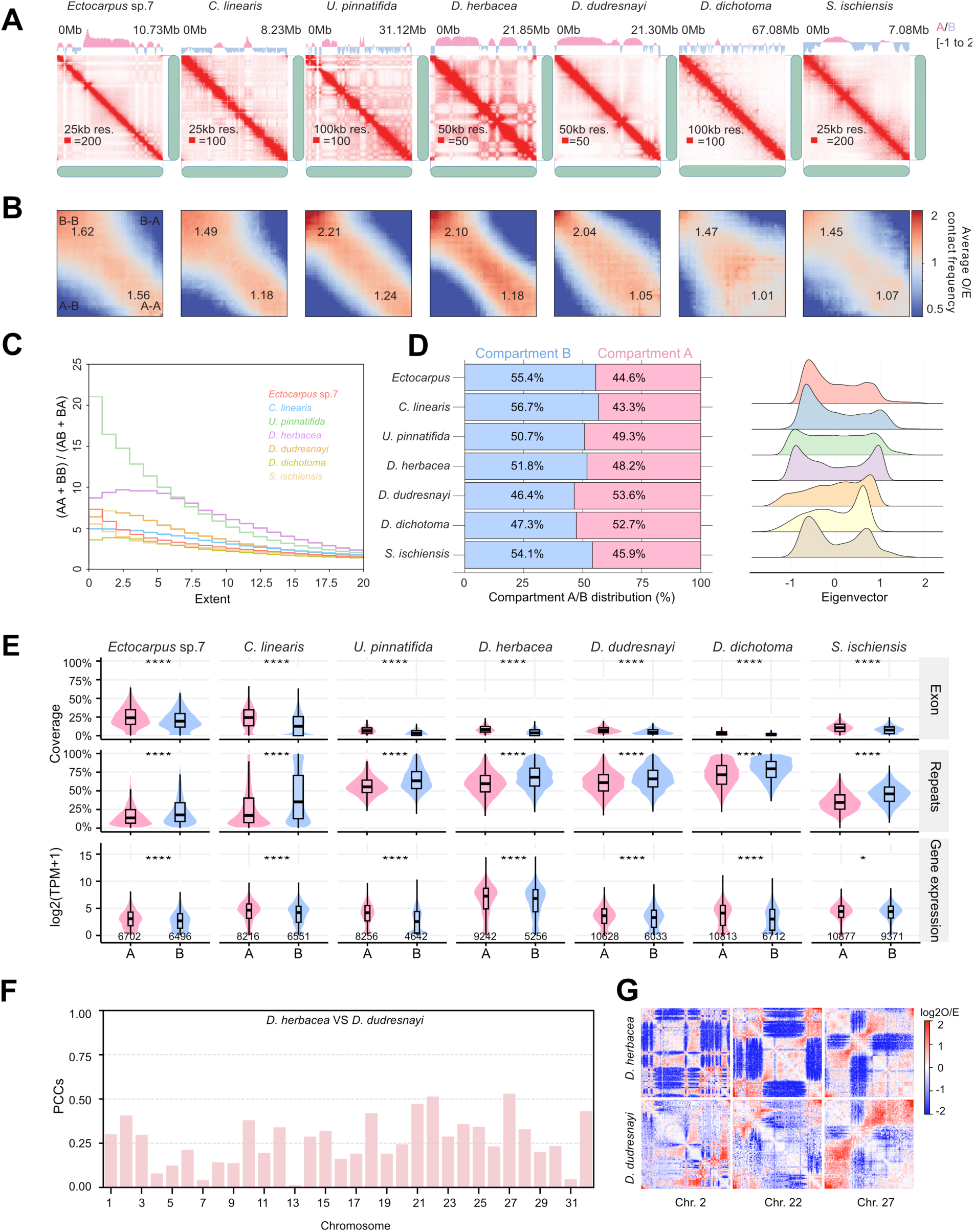
Comparison of chromosome conformation in brown algae. (**A**) Chromosome 1 of each brown algal species with genome compartmentalization. Eigenvector values for each chromosome are shown above the corresponding Hi-C matrix in juicebox^45^. (**B**) Saddle plots of genome-wide interactions showing compartment contact interactions in cis for each species (numbers are relative interactions for AA and BB interactions type). (**C**) Saddle strength quantifying compartmentalization interaction frequencies. The metric represents the ratio of intra-compartment (A–A and B–B) to inter-compartment (AB or BA) interactions, normalized by expected contact frequencies. Genomic bins are sorted according to eigenvector values obtained from Hi-C eigenvector analysis. The ‘extent’ parameter defines the range of eigenvector quantiles included in the calculation: an extent of 0 considers only the most extreme A and B bins, thereby capturing the strongest compartmentalization signal, whereas increasing extent values progressively incorporate bins closer to the center of the eigenvector distribution, where compartment identity is weaker. (**D**) Genomic distribution of chromatin compartments. Bars show the proportion of each genome assigned to A compartments (gene-rich, transcriptionally active) or B compartments (gene-poor, heterochromatic) based on Hi-C eigenvector analysis. (**E**) Violin plot shows the percentage of exons, repeats, and gene expression in each compartment across species. Numbers below brackets represent the number of genes. The lower and upper hinges of the box correspond to the first and third quartiles (the 25th and 75th percentiles). The upper whisker extends from the hinge to the largest and smallest values no further than 1.5x IQR from the hinge (Inter-Quartile Range, distance between the first and third quartiles). Significance was determined by a two-sample Wilcoxon rank sum test (****: p < 0.0001, *: p <=0.05). (**F**) Pearson coefficient correlations (PCCs) of linearly transformed Hi-C contact matrices between homologous chromosomes of *D. herbacea* and *D. dudresnayi* across all chromosomes; (**G**) Examples show strongly correlated orthologous chromosomes between *D. herbacea* and *D. dudresnayi* at 50k resolution.

To assess whether differences in compartment strength between species reflect broader changes in compartmental organization, we analyzed the proportion of A and B compartments and the distribution of eigenvector values across species. Given that larger genomes typically harbor more TEs^25,26^, which are often associated with heterochromatin and B compartments, a higher prevalence of B compartments in species with larger genomes is expected. However, our analysis revealed strikingly similar A/B compartment fractions across all species examined, with no clear correlation between genome size and compartment proportions (**Fig. 2D**). Therefore, both the organization and relative abundance of A/B compartments are largely independent of genome size, even though their strength may vary. Note that consistent with the open/closed nature of A/B compartments, compartment A has more exonic sequence and compartment B has more repeats, and genes located within compartment A exhibited significantly higher expression levels than those in compartment B across all species analyzed (**Fig. 2E**).

Previous work reported that the *Ectocarpus* genome lacks small-scale chromatin structures such as TADs^31^. Unexpectedly, visual inspection of Hi-C maps revealed domain-like structures in *U. pinnatifida*, *D. herbacea* and *D. dichotoma* (**Fig. S8A**). These domains, which exhibit strong self-interaction frequencies and are delimited by pronounced insulation boundaries, ranged in size from 258.2 kb in *D. dudresnayi* to 355.1 kb in *U. pinnatifida* (**Fig. S9**). To further characterize these structures, we generated insulation profiles using multiple distance ranges (50-500 kbp, **Fig. S8B**). Based on visual inspection, cis-chromatin contacts within 100 kbp provided optimal boundary resolution and were used for subsequent analysis, leading to the identification of insulated regions that we named ‘boundaries’. To assess whether genome size relates to chromatin organization, we compared the mean insulation strength of boundaries with genome size and observed a moderate positive correlation (Pearson’s r = 0.53), although this association was not statistically significant (p = 0.223), likely due to the small sample size (**Fig. S8C**). No enrichment of genomic features, including TEs, exons, or introns, was observed at these boundaries (**Fig. S8D**). Therefore, although morphologically complex species with large genomes tended to exhibit domain-like structures, these could not be linked to any specific genomic feature.

In the outgroup species *S. ischiensis*, the Hi-C contact map revealed plaid patterns indicative of alternating open and closed chromatin regions, as well as centromere interaction clusters and centromere aggregations (**Fig. S6**). No self-interacting chromatin domains were detected, even at higher resolutions (**Fig. S8A**). However, we observed prominent inter-chromosomal interaction clusters involving *S. ischiensis* chromosome 5. These regions are enriched for LINE retrotransposons (see **Fig. S10**).

Brown algae exhibit a relatively conserved chromosome number, ranging from 27 to 32. Macro- synteny analyses reveal largely 1:1 orthology across most chromosomes when compared to out- group species (**Fig. 1C**). Given this conservation, we asked whether it extends to 3D chromatin architecture. Hi-C matrices were visually inspected in Juicebox^45^ to assess whether orthologous chromosomes could be distinguished based on contact patterns. Notably, *D. herbacea* and *D. dudresnayi*, which diverged ∼11 Mya, display clear 1:1 synteny, composed of a few large syntenic blocks (see **Fig. 1C**), and their orthologous chromosomes share remarkably similar inter-chromosomal interactions (**Fig. S6F-G**). To quantify this similarity at the 3D chromatin level, Hi-C contact matrices were normalized to uniform 50 kb bins. Pearson correlation coefficients (PCCs) across the 32 orthologous chromosomes ranged from 0.012 to 0.53 (**Fig. 2F**), with higher PCCs reflecting substantial conservation of chromatin organization (**Fig. 2G**). By contrast, comparisons between other pairs of sister species such as *Ectocarpus* and *C. linearis* showed lower PCCs (**Table S6**), likely due to greater divergence times (110 My). These results indicate that chromatin conformation is largely conserved for at least ∼11 million years of brown algal evolution, but not over larger evolutionary timescales.

In summary, A/B compartmentalization and its functional associations with gene density, repeats, and gene expression are largely conserved across 450 My of evolution encompassing brown algae and the outgroup *S. ischiensis*. By contrast, domain-like structures vary: species with greater morphological complexity show stronger, more distinct compartmentalization, whereas simpler species, including the outgroup, exhibit weaker domains.

### Unique chromatin interaction patterns of sex chromosomes or sex-homologs

Like many early-diverged plants^23,46^, sex determination in most brown algae occurs in the haploid stage of the life cycle, and haploid individuals of dioicous species possess either female (U) or male (V) sex chromosomes^23,29^. Previous analyses have shown that U/V sex chromosomes exhibit unusual linear genomic features, including distinct repeat content, gene density, and levels of sequence divergence compared with autosomes^25,47^. To explore whether these distinctive linear properties are reflected in higher-order chromatin organization, we analyzed the 3D topology of the U and V chromosomes across our sampled species. We examined contact frequencies, domain formation, and compartmentalization patterns, aiming to determine whether sex chromosomes adopt unique spatial conformations that could relate to their specialized genomic architecture and evolutionary dynamics.

Intriguingly, we observed prominent inter-chromosomal interactions between the sex-determining region (SDR) and multiple autosomes in *D. dichotoma*, *D. herbacea*, and *D. dudresnayi* (**Fig. 3A, Figs. S6, S11A**), suggesting a lineage-specific propensity for the sex chromosome to engage in trans-contacts, and this is not constrained by chromosome size (**Fig. S12A-B**). Note that these interactions do not result from self-interactions of tandem repeats (**Fig. S11B**) or mapping artifacts (**Fig. S11E**), but instead represent *bona fide* preferential contacts. To comprehensively investigate whether interacting regions share genomic features, we profiled the distribution of TE families, exons, compartment A/B regions, and centromeric sequences genome-wide for *D. dichotoma* (**Fig. S11F**). This analysis indicated that chromatin interactions are not primarily driven by these specific genomic features, indicating that alternative organizational principles may govern 3D genome structure in sex chromosomes.

**Fig. 3.**
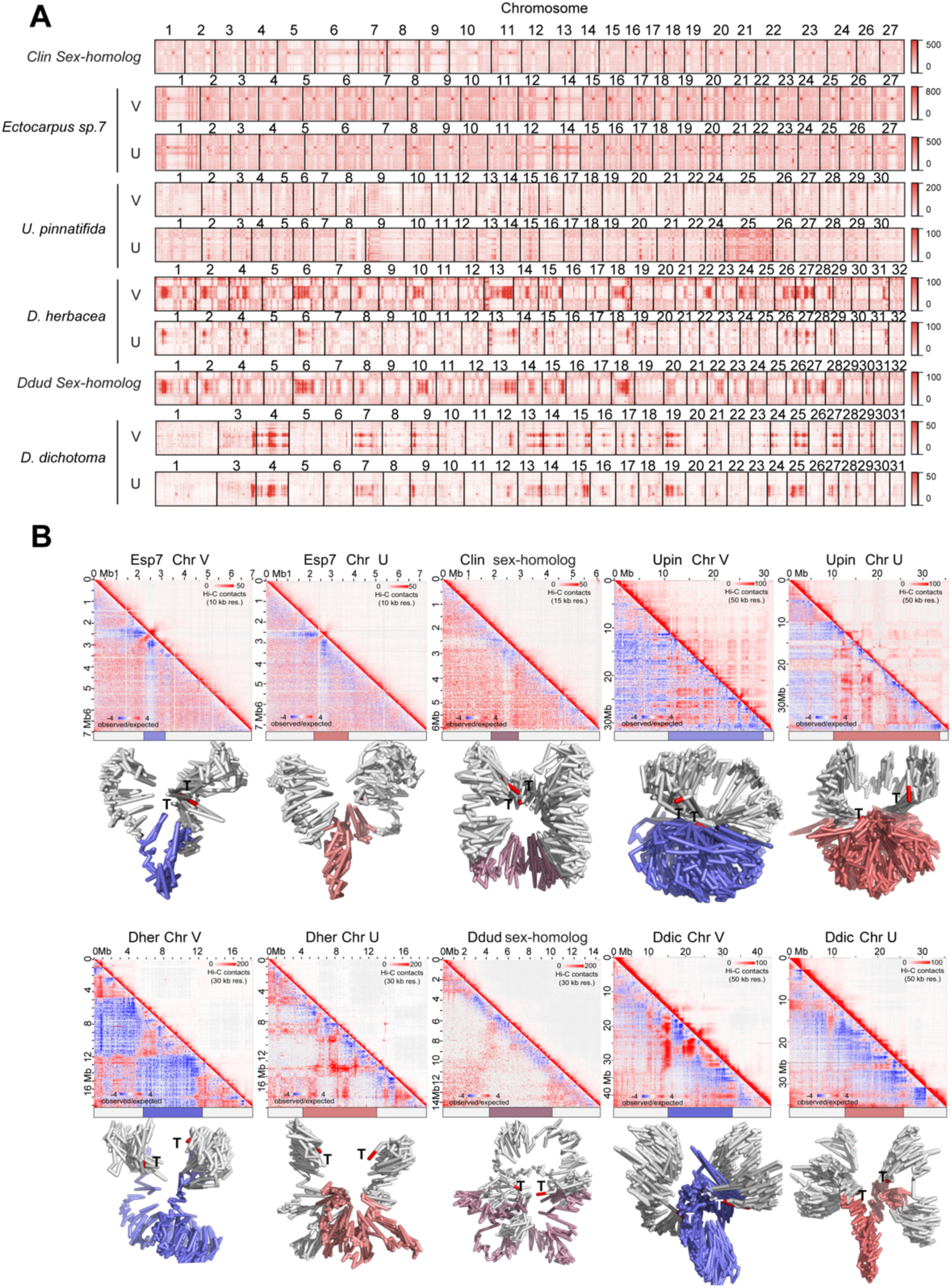
3D chromatin configuration of sex chromosome or sex-homologs in brown algae. (A) Hi-C contact maps showing inter-chromosomal interactions between the sex chromosome (or sex homolog) and autosomes for each species; (B) Hi-C map and reconstructed 3D configurations of sex chromosomes at 25 k resolution Hi-C data, maximum likelihood approach was employed to construct 3D structures from Hi-C data of *Ectocarpus* sp.7^31^, *C. linearis*, *U. pinnatifida*, *D. herbacea*, *D. dudresnayi* and *D. dichotoma* with a default setting in 3DMax^49^. SDRs or SDR-homologs in the V, U chromosomes and sex-homolog are colored in slate, deepsalmon and purple, respectively, and telomeres are colored in red, pseudoautosomal regions (PARs) are colored in grey.

Sex chromosomes were predominantly assigned to the B compartment across species (**Fig. S13A-C**), while sex-determining regions (SDRs) show enrichment in compartment A. Unexpectedly, repeat elements are more enriched in compartment A than compartment B on sex chromosomes, inverting the typical genome-wide pattern where compartment A is gene-rich and compartment B is repeat-rich (**Fig. S13D**). Gene expression patterns in A versus B compartments vary across species, with significantly higher expression in A compartment observed only in *Ectocarpus* (**Fig. S13E**).

All examined SDRs exhibit strong insulation, indicating this is a conserved feature of brown algal sex chromosome organization (**Fig. 3B**). Notably, *D. herbacea* shows sex-specific differences in chromatin architecture: the female SDR is highly organized into smaller, self-interacting chromatin domains, while the male SDR lacks this organization (**Fig. 3B**). Genes located at chromatin domain boundaries displayed elevated expression compared to those within or outside domains on sex chromosomes (**Fig. S14**). Together, these data demonstrate that brown algal sex chromosomes have evolved conserved yet specialized chromatin architectures featuring enhanced insulation and reorganized domain structures, establishing 3D genome organization as a fundamental mechanism underlying sex chromosome evolution and regulation.

We next asked whether sex chromosome chromatin architectures persist following evolutionary transitions to hermaphroditism, where former sex chromosomes become autosomes^48^. Our dataset captures two independent transitions: *Ectocarpus* versus *C. linearis*, and *D. herbacea* versus *D. dudresnayi*, with the latter representing a recent transition occurring within the last 11 million years. Strikingly, in both lineages, the chromatin organization of former sex chromosomes ("sex-homologs") has retained the architectural signature of the ancestral male (but not female) sex chromosome rather than adopting autosomal patterns (**Fig. S15**). This evolutionary constraint is most pronounced in *D. herbacea*, where the large SDR and recent transition timeline provide a clear window into chromatin evolution (**Fig. 3B**). These findings demonstrate that three-dimensional chromatin organization exhibits remarkable evolutionary inertia, with former sex chromosomes maintaining male-specific architectural features long after their functional role in sex determination has been lost.

### Longterm co-evolution of centrophilic retrotransposons and brown algal centromeres

The structures of regional centromeres exhibit extreme variation among eukaryotes and typically evolve rapidly^50,51^. The centromeres of many species feature centrophilic retrotransposons, specific TE families that target the centromere and in some cases directly correspond to the sequence occupied by the centromeric histone variant CENH3^52^. Centrophilic retrotransposons can co-occur with centromeric satellite repeats, forming a spectrum from a relatively minor contribution to centromeric sequence, e.g., *ATHILA* elements in *Arabidopsis thaliana*^53^, to dominating the centromeres of many chromosomes, e.g. the *CRM* elements of maize^54^. In other species, centrophilic retrotransposons solely define and constitute the centromeres, e.g. *Bryco* elements in the moss *Physcomitrium patens*^55^ or *ZeppL* elements in the green alga *Chlamydomonas reinhardtii*^56,57^.

We previously identified centromeric regions in *Ectocarpus* sp. 7 based on strong centromere-centromere interactions in inter-chromosomal Hi-C contact maps^31^. Although these interaction clusters span hundreds of kilobases, we mapped the putative centromeres to highly localized regions that typically lack satellite DNA but are defined by the presence of two specific LTR retrotransposon families from the *Metaviridae*/Ty3 group, termed *ECR* (*Ectocarpus Centromeric Retrotransposon*) elements. **Fig. 4A** shows a representative pair of chromosomes from *Ectocarpus* sp. 7, with a single broad region of strong centromere-centromere interaction clusters visible on inter-chromosomal Hi-C matrix in Juicebox^45^, in addition to prominent telomere-telomere interaction clusters. The two centromeric regions harbor short (∼30-40 kb) clusters of the *ECR-1* (orange) and *ECR-2* (vermillion) elements, and with no other repeats consistently present these retrotransposons presumably correspond to the epigenetic centromere (i.e., the region featuring CENH3). While many *ECR-1* elements are intact and potentially functional, all *ECR-2* copies are degraded and the family may be in the process of extinction^31^.

**Fig. 4.**
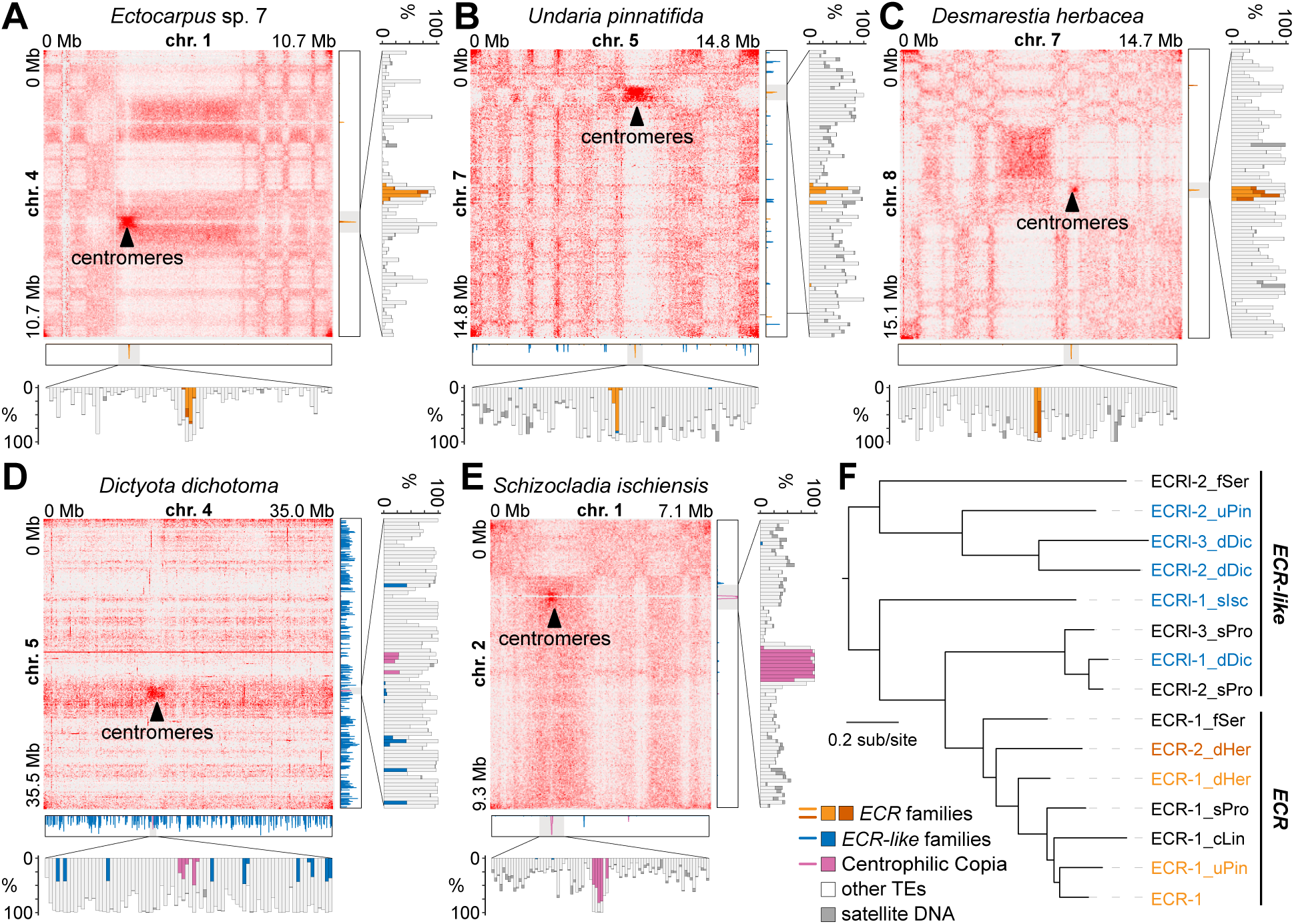
Centromere profiles across brown algal genomes. Representative examples show centromere clustering between two chromosomes in Hi-C interaction frequency maps, along with the distribution of centromeric retrotransposons (*ECR* and Copia), non-centrophilic *ECR-like* elements, other TEs and satellite DNA in 10 kb windows for (A) *Ectocarpus* sp. 7, (B) *U. pinnatifida*, (C) *D. herbacea*, (D) *D. dichotoma*, and (E) *S. ischiensis*. Chromosome-scale plot shows only centromeric retrotransposons and *ECR*-like elements, centromere-centered panel shows all repeat types. (F) Maximum likelihood phylogeny of *ECR* and *ECR-like* elements constructed from Gag and Pol protein sequences under the LG+F+I+G4 model. All *ECR* elements form genomic clusters, and those forming specific centromeric clusters in panels A-E are colored orange or vermillion. All nodes received > 95% ultrafast bootstrap support with the exception of those connecting *ECR-1*, *ECR-1_uPin* and *ECR-1_cLin*.

We observed prominent centromere-centromere and telomere-telomere interaction clusters in the Hi-C contact maps of all species (**Fig. 4B-E, S6**). This organization appears to be common to both brown algae and the outgroup *S. ischiensis*, and genome folding on a broad chromosomal level may be consistent across the whole lineage. We searched for homologs of *ECR-1* in each genome and asked whether these elements also form clusters within the broad centromeric regions defined from the Hi-C maps. In *U. pinnatifida*, we identified two homologous families: *ECR-1_uPin* mostly (but not exclusively) forms short centromere-localized clusters (**Fig. 4B and S16C,** orange), whereas *ECRl-2_uPin* exhibits a genome-wide distribution (**Fig. 4B and S16C,** blue). In *D. herbacea*, the families *ECR-1_dHer* and *ECR-2_dHer* both co-occur in short centromeric clusters (**Fig. 4C, S16D**). Although Hi-C contact maps are unavailable, we also identified *ECR* families that either form highly localized clusters or have genome-wide distributions in the contiguous genome assemblies of the species *Scytosiphon promiscuus* and *Fucus serratus*^25^ (**Fig. S16B and S16F**), mirroring the pattern observed in *U. pinnatifida*. In *C. linearis* the situation is less clear; we identified a single *ECR* family that is localized within the Hi-C defined centromeric regions (**Fig. S16**A, **S6A**), however it is only present on 10 of the chromosomes and is frequently fragmented, suggesting that in may be in the process of elimination from the genome. At least two other repeats are enriched within the broadly defined centromeric regions, although these resemble neither TEs nor satellites and their role (if any) in centromere function is unclear. One appears to be a multicopy gene encoding a protein with a predicted P-loop domain (Pfam PF07999).

Conversely, in *D. dichotoma* we identified three abundant *ECR* families that all exhibit genome-wide distributions (**Fig. 4D and S16E**). Similarly, in *S. ischiensis* the sole *ECR* family does not exhibit centromeric clustering and is present at low copy numbers elsewhere in the genome (**Fig. 4E and S16G**).

We performed a phylogenetic analysis using the combined Gag and Pol protein sequences of all identified *ECR* elements. Interestingly, the centromere-associated *ECR* elements form a distinct and robustly supported sub-lineage (ultrafast bootstrap value 100) within the wider diversity of *ECR* elements (**Fig. 4F**). Thus, the lineage can be divided to genuine centrophilic *ECR* elements and non-centrophilic *ECR-like* elements (**Fig. 4F**), which can co-occur in a single genome (*U. pinnatifida, S. promiscuus* and *F. serratus*), be present only as centrophilic *ECR* elements (*Ectocarpus* sp. 7 and *D. herbacea*), or only as *ECR-like* elements (*D. dichotoma* and *S. ischiensis*). Thus, it appears that centrophily evolved from a non-centrophilic LTR retrotransposon and had emerged in the common ancestor of the brown algal crown radiation (BACR) at least 160 MY^22^.

We next searched for any repeats that are enriched in the broadly defined centromeric regions of *D. dichotoma* and *S. ischiensis*. Surprisingly, we identified single families of Copia (*Pseudoviridate/*Ty1) LTR retrotransposons that form discrete clusters in the two species (**Fig. 4D, E**). Phylogenetic analysis of all Copia elements identified in the six analyzed species revealed that these putatively centrophilic elements do belong to the same weakly supported lineage (**Fig. S16H**). However, this lineage also includes three other Copia family from *S. ischiensis* that have a genome-wide distribution, and families from *Ectocarpus* sp. 7, *D. herbaceae* and *C. linearis* that are present genome-wide. Thus, it is possible that there have been two independent transitions to centrophily during the evolution of *D. dichotoma* and *S. ischiensis*, or alternatively, the centrophilic Copia elements could represent an ancestral state with multiple transitions to centrophobic insertion patterns.

Despite featuring distinct LTR retrotransposons, there are parallels between the putative centromere structures of most brown algae and *S. ischiensis*. The LTR clusters are generally short, typically spanning tens of kilobases as opposed to hundreds of kilobases as common in other species^58^. This appears to be true regardless of the repeat content of the genome, for example, in the more repeat-rich genomes of *U. pinnatifida, D. herbacea* and *D. dichotoma*, the centromeres do not exhibit an elevated repeat content relative to the surrounding sequence (**Fig. 4B-D**). As described for *Ectocarpus* sp. 7^31^, satellite DNA is also conspicuously absent from most centromeres in all species (**Fig. 4A-E**), reinforcing the interpretation that the centrophilic retrotransposons (either *ECR* or Copia) may directly correspond to the epigenetic centromeres across brown algae.

### The ancestral brown algal karyotype and accelerated chromosome evolution in the Ectocarpales

Chromosome evolution involves major genome rearrangements that occur within (e.g., inversions) or between chromosomes, with the latter category including several types of chromosome fusion and fission, in addition to reciprocal translocations. As introduced above, chromosome number ranges from 27 to 32 in our sample of genomes, and the availability of chromosome-level assemblies enables us to reconstruct chromosome evolution over ∼240 million years of brown algal evolution for the first time. As presented in **Fig. 1C**, one-to-one chromosome-scale synteny is evident among the majority of chromosomes (n=21) in a comparison of *D. dichotoma, D. herbacea* and *D. dudresnayi*, and *U. pinnatifida*. This number increases to 26 chromosomes in a pairwise comparison of *D. dichotoma* (here the outgroup) and *D. herbacea*, which appear to be differentiated by only three inter-chromosomal rearrangements. Although the order of these events cannot be determined without additional outgroups, if we arbitrarily assume that *D. dichotoma* represents the ancestral state then the rearrangements can be expressed as: i) fission of *D. dichotoma* chromosome 6, giving rise to *D. herbacea* chromosomes 29 and 30, ii) fission of *D. dichotoma* chromosome 1 and subsequent fission of one fragment to chromosome 12, giving rise to *D. herbacea* chromosome 30, and iii) reciprocal translocation between *D. dichotoma* chromosomes 27 and 31, yielding *D. herbacea* chromosomes 21 and 24 (see **Fig. 1C**). Furthermore, based on parsimony we can infer that the ancestral brown alga had either 31 or 32 chromosomes, with the point of distinction our inability to determine if *D. dichotoma* chromosome 6 is the result of a fusion, or if *D. herbacea* chromosomes 29 and 30 are the product of a fission.

The remaining inter-chromosomal rearrangements can be polarized relative to this ancestral karyotype. *D. dudresneyi* features one unique rearrangement event, the fission of *D. herbacea* chromosome 30 and subsequent fusion of a fragment to chromosome 19, which can be inferred to have occurred in the last 11 million years. As with the inferred fission+fusion event between *D. herbacea* and *D. dichotoma*, it is also possible that this event was instead a reciprocal translocation that occurred close to the telomere on one of the chromosomes, leaving no trace in the synteny analysis. The *U. pinnatifida* chromosomes 1 and 25 are both products of fusion events that are unique to this species, whereas chromosomes 12 and 29 result from a reciprocal translocation that is also shared by *Ectocarpus* sp. 7 and *C. linearis*, suggesting it occurred prior to the common ancestor of Ectocarpales and Laminariales. Strikingly, the Ectocarpales feature the most derived karyotypes, especially *C. linearis* (**Fig. 1C**). Although the complexity of several of these rearrangements complicates inference, we estimate that two fusions occurred in the ancestor of Ectocarpales, with two further fusions and a reciprocal translocation occurring on the lineage leading to *Ectocarpus* sp. 7, and two distinct fusions and seven reciprocal translocations occurring on the lineage leading to *C. linearis* (**Fig. 5A, S17**). Despite the higher rate of rearrangements in the Ectocarpales, 8 chromosomes exhibit a 1-1 relationship across all analyzed brown algal genomes, including the sex (or sex-homolog) chromosome.

**Fig. 5.**
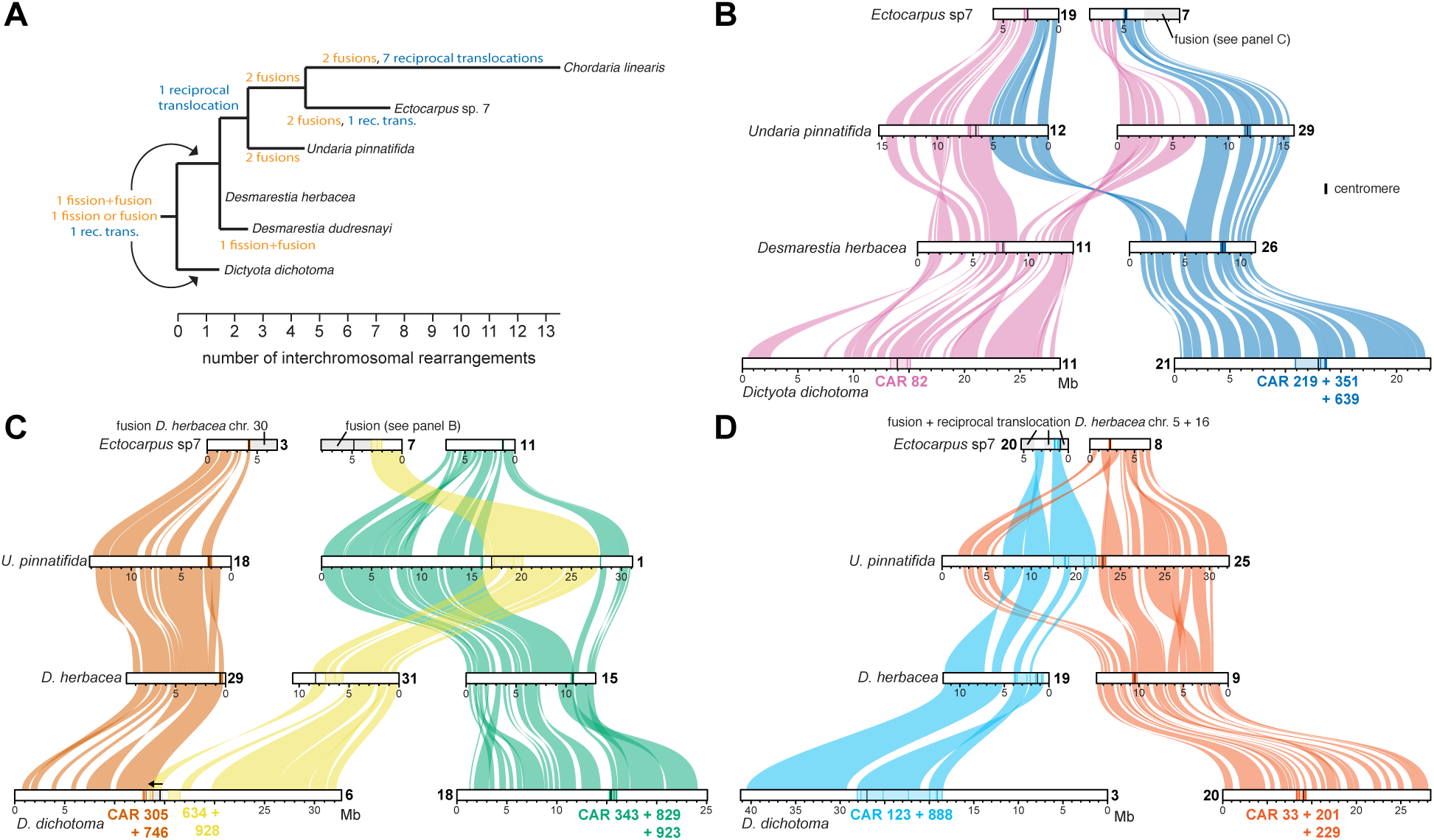
Inter-chromosomal rearrangements in brown algal genomes. A) Phylogeny of analyzed brown algal genomes where branch lengths correspond to the number of inter-chromosomal rearrangements that occurred between each node. B-D) Syntenic relationships among a representative selection of chromosomes associated with inter-chromosomal rearrangements, showing a reciprocal translocation (B), a putative Robertsonian translocation (C) and nested fusions (C, D). Chromosomes are colored relative to *D. herbacea* and contiguous ancestral regions (CARs) defined by AGORA^49^ that flank centromeres are highlighted (each line corresponds to a gene present in the CAR). A predicted centromeric inversion on *D. dichotoma* chromosome 6 (panel C) is represented by a black arrow. For all Ectocarpales rearrangements see **Fig. S17**.

Centromeres are frequently involved in chromosomal rearrangements, with centromere-proximal rearrangement breakpoints associated with end-to-end fusions (including Robertsonian translocations) and nested fusions (i.e., centric insertion)^17,59^. To further address the origin of brown algal chromosomal rearrangements, we mapped centromere locations on to a representative selection of rearrangements (**Fig. 5A-C**). To determine if centromere locations are conserved among species, we performed an ancestral genome reconstruction using AGORA, which assembles contiguous ancestral regions (CARs) featuring multiple genes inferred to have been adjacent in an ancestral genome^60^. This analysis revealed that centromeres are generally located within the same CAR, or between the same adjacent CARs, in each genome, suggested ancient conservation of centromeric locations across brown algae (**Fig. 5A-C**). This is true even for *D. dichotoma*, despite the difference in the identity of centrophilic LTR retrotransposons in this species (see **Fig. 4D**).

Most reciprocal translocations were not associated with centromeres, as exemplified by the reciprocal translocation common to *U. pinnatifida* and *Ectocarpus* sp. 7 (**Fig. 5A, B**). Conversely, many of the fusions (or fissions) potentially involved centromeres. The *D. herbacea* chromosomes 29 and 31 are both acrocentric, suggesting that the fusion or fission event that differentiates them from *D. dichotoma* could have been a Robertsonian translocation (**Fig. 5C**). However, at least one centromeric inversion has likely occurred since this event, complicating inference. *U. pinnatifida* chromosome 1 appears to have resulted from centric insertion of the acrocentric *D. herbacea* chromosome 31 into the centromere of *D. herbacea* chromosome 15 (**Fig. 5C**). Similarly, *U. pinnatifida* chromosome 25 is the product of a nested insertion of *D. herbacea* chromosome 19 into 9, although in this case the insertion point appears to have been adjacent to the centromere rather than within it, with the ancestral chromosome 19 centromere having been lost in the fused chromosome (**Fig. 5D**). Note that *Ectocarpus* sp. 7 chromosome 7 is the product of an end-to-end fusion between the acrocentric *D. herbacea* chromosome 31 and the metacentric *U. pinnatifida* chromosome 29, which resulted in the loss of the chromosome 31 centromere (**Fig. 5C, D**). Many of the other rearrangements in Ectocarpales are also adjacent to a centromere on at least one of the chromosomes (**Fig. S17**).

Finally, we used the ancestral brown algal genome reconstruction to investigate syntenic relationships with *S. ischiensis*, which last shared a common ancestor with brown algae ∼450 mya (see **Fig. 1B**). Arbitrarily assuming that *D. herbacea* represents the ancestral state of brown algal chromosomes, we painted the *S. ischiensis* chromosomes based on the orthology relationships of genes between the two species. Essentially all of the *S. ischiensis* chromosomes feature a mixture of genes from different brown algal ancestral chromosomes (**Fig. S18**), with the possible exceptions of chromosomes 24 and 26 that correspond to *D. herbacea* chromosomes 17 and 26, respectively, with only limited mixing with other chromosomes. This analysis shows that *S. ischiensis* shares low macrosynteny with brown algal genomes, confirming a result derived previously from a more fragmented version of the genome assembly^25^.

Overall, our results reveal that the ancestral brown alga likely had 31 or 32 chromosomes, with a small number of centromere-associated fusion/fissions and mostly non-centromeric reciprocal translocations differentiating the karyotypes of most species. The rate of chromosome evolution appears to have accelerated in the Ectocarpales, and especially in *C. linearis*, which exhibits the most derived brown algal karyotype resulting from extensive reciprocal translocation in this lineage (**Fig. 5A**).

## Discussion

### Conservation and variation in 3D chromatin architecture across evolutionary scales

Our analyses reveal a striking conservation of higher-order chromatin architecture despite wide variation in genome size and repetitive element content. These results suggest that key features of 3D genome folding: such as chromosomal territories, centromere clustering, telomere clustering, and A/B compartments are maintained under strong evolutionary constraint across the brown algal lineage, mirroring patterns in animals^3^, plants^61^, and fungi^62^, and reflecting a conserved functional architecture for gene regulation and chromatin organization. The presence of A/B compartmentalization in brown algae, which diverged from plants and animals over a billion years ago^1^, indicates either deep ancestral origins or convergent evolution driven of this feature by fundamental nuclear constraints. Critically, despite substantial variation in brown algal genome sizes neither eigenvector distributions nor compartmentalization strength correlate with genome size. This independence from genome composition suggests that spatial organization may serve essential regulatory functions that go beyond genomic complexity.

In animal lineages, chromatin domains are defined by local insulation boundaries and shaped by architectural proteins such as CTCF, cohesin, and condensin^1^. In contrast, both plants^36,61^ and brown algae lack canonical insulator proteins like CTCF, and there are no prominent TADs as commonly observed in animals^63^. In plants with large genomes, however, domains can be found and their boundaries are often associated with transcriptionally active regions specific histone modifications^10,64^ (e.g., H3K4me3 or H3K9me2) or chromatin state transitions, rather than fixed sequence motifs. Similarly, in brown algae, non-canonical chromatin domains are detectable in some species with larger genomes, exhibiting elevated insulation levels that suggest alternative, genome size or TE content-dependent mechanisms for domain formation. Notably, the relationship between gene expression and chromatin domains appears inconsistent across brown algal species, unlike in animals where genes near domain boundaries often exhibit elevated expression^1^. We found no evidence for enrichment of specific genomic features such as gene families or TE classes at insulation boundaries. This contrasts with other eukaryotic systems, where boundary elements often coincide with active genes, tRNAs, or specific DNA-binding motifs^34,65^, though CTCF involvement varies and boundary mechanisms can differ among lineages. In brown algae, boundary positioning may instead be influenced by other, as yet unidentified factors, such as nucleosome organization^66^, histone modifications^1^, or structural RNAs^67^. Alternatively, boundaries may form passively as a consequence of chromatin state transitions or replication timing domains, as proposed in some plant species^62,68^.

Taken together, our results demonstrate that A/B compartmentalization is a conserved and robust feature of brown algal nuclear organization, maintained over evolution despite substantial differences in genome size and repeat content. In contrast, chromatin domains defined by local insulation minima are observed only in larger genome species, suggesting that domain-level genome folding is not a universal characteristic within this lineage. The lack of correlation between genome size and insulation strength contrasts with models in other eukaryotes where genome expansion and TE load are major drivers of architectural complexity. Our findings thus support the concept that 3D genome architecture is shaped by lineage-specific evolutionary constraints^44^ rather than being universally determined by genome size alone.

### Conserved macrosynteny and chromosomal stability over evolutionary time

We generated high-quality, chromosome-level genome assemblies for five brown algal species and one outgroup. These assemblies markedly improve completeness and contiguity over previous versions, enabling detailed analyses of genome evolution and 3D chromatin organization in this understudied eukaryotic lineage.

Ancestral reconstruction of chromosomes or linkage groups has emerged as a powerful analytical technique in many eukaryotic groups, revealing various degrees of conserved macrosynteny among mammals^27^, vertebrates^69^, and even some^17^, but far from all^70^, bilaterians. The concepts of macrosynteny and ancestral linkage groups have long been associated with the Muller elements of drosophilids, the gene content of which remains largely conserved over more than 50 million years of evolution despite the gene order itself being frequently rearranged between species by inversions^71^. Similar ancestral linkage groups have been defined for lepidopterans (Merian elements)^72^ and rhabditid nematodes (Nigon elements)^73^, as well as in some plant groups such as the Brassicaceae^74^.

Our chromosome-level assemblies revealed comparably extensive macrosynteny across brown algae, enabling an ancestral karyotype of either 31 or 32 linkage groups to be constructed. These ancestral chromosomes had become established at least 240 million years ago, with earlier evolutionary relationships made uncertain by the extensive breakdown of macrosynteny between brown algae and the outgroup *S. ischiensis*. Gene collinearity (i.e., synteny) is also extensive among brown algal genomes (Fig. 1C, 5B-D), although some intrachromosomal rearrangements between species have occurred as in other eukaryotic groups with conserved macrosynteny. Notably, chromosomes that have arisen via fusion generally retain their ancestral macrosynteny patterns (Fig. 5C, D), with limited intrachromosomal rearrangements between the different ancestral linkage groups. While these may simply represent very recent events, similar patterns are observed in the Muller and Merian elements of insects^71,72^, and it may be the case that macrosynteny is selectively retained to maintain *cis*-regulation between genes^27,72^. Overall, as in other complex eukaryotic lineages, conserved and ancient macrosynteny may reflect strong evolutionary constraints contributing to long-term genomic and adaptive stability.

Nevertheless, we detected substantial chromosomal rearrangements in the Ectocarpales, corresponding to elevated rates of both fusions and reciprocal translocations. Interestingly, this group comprises species with the smallest and most compact genomes. Such compact genomes, typically characterized by higher gene density, are often linked to elevated recombination rates^75^ which can in turn promote chromosomal rearrangements. Smaller chromosomes have also been more substantially involved in fusions in vertebrate^76^ and lepidopteran^72^ evolution. Moreover, Ectocarpales species have short life cycles^21^ with frequent meiotic divisions, a feature that may further enhance recombination activity and thus accelerate the rate of chromosomal reorganization^77^ observed in this lineage. Finally, the Ectocarpales also exhibit reduced developmental complexity relative to the other analyzed brown algae and although speculative we cannot rule out that this evolutionary transition to simpler morphologies has resulted in relaxed constraint on macrosynteny.

### Distinct chromatin features of sex chromosomes and sex-homologs

Little is known about the 3D chromatin organization of sex chromosomes across eukaryotes^20,78^. Sex chromosomes in brown algae exhibit distinct three-dimensional architectures that set them apart from autosomes. Although largely assigned to the transcriptionally inactive B compartment, their non-recombining SDRs consistently localize to the active A compartment. This organization is further marked by an inversion of genomic feature distribution, with repeat elements enriched in active A compartments. It is possible that this reflects the conspicuous presence of TEs in SDR gene introns^25,47,40^. Strong insulation of all examined SDRs points to a conserved structural hallmark of non-recombining regions across lineages. Evolutionary analyses further reveal that former sex chromosomes retain male-, but not female-, specific chromatin signatures long after transitions to hermaphroditism, underscoring the persistence of ancestral features^25,79^. In brown algae, hermaphrodites are derived from male lineages that acquired female-specific genes^25,48^. Accordingly, our results suggest that V-specific chromatin architecture is maintained through this transition. Together, these results show that chromatin organization is both a defining feature of sex chromosomes and an evolutionarily constrained trait that endures beyond its original functional role.

### Conserved centromeric locations and long-term co-evolution with centrophilic retrotransposons

The extensive centromere-centromere contacts in the Hi-C maps of brown algae and *S. ischiensis* enabled us to investigate centromere structure over 450 million years of evolution. As we recently established in *Ectocarpus* sp. 7^31^, the centromeres of all species feature a short cluster of LTR retrotransposons that belong to specific families that are conspicuously absent from the rest of the genome. These centrophilic LTR families have presumably evolved to target the centromere of their respective genome, as has been established for several distinct groups of retrotransposons in specific plant and animal genomes^52,80^. The absence of satellite DNA or any other consistent repetitive sequences also suggests that the centrophilic retrotransposon clusters likely correspond to the epigenetic centromere, as in *P. patens*^44^ and *C. reinhardtii*^57^. Antibodies targeting CENH3, which are currently unavailable for brown algae, will be required to test the association between the centrophilic LTR elements and the epigenetic centromere.

Our analyses also revealed the long-term co-evolution between *ECR* LTR retrotransposons and the centromeres of a subset of species that correspond to the brown algal crown radiation (BACR). *ECR-1*, the intact and presumably active centrophilic retrotransposon of *Ectocarpus* sp. 7^31^, forms a distinct evolutionary lineage relative to all other *Metaviridae/*Ty3 families in the *Ectocarpus* sp. 7 genome^31^. We identified several other members of this lineage across the brown algal genomes, including both centrophilic and non-centrophilic families. Strikingly, the centrophilic elements, here referred to as genuine *ECR* elements, form a clade within the wider diversity of non-centrophilic *ECR-like* elements, suggesting an evolutionary transition to centrophily that occurred at least 160 mya in the common ancestor of the BACR. Although their long-term persistence in the genome of *C. linearis* is unclear, the centrophilic *ECR* elements appear to be the core centromeric components of all other BACR genomes analyzed, and searches for *ECR* elements are likely to prove useful for centromere identification across the majority of brown algal species.

Such long-term co-evolution between a specific lineage of retrotransposons and centromeres is most closely paralleled by the *CRM* elements that are present at the centromeres of many angiosperm species. *CRM* elements represent a centrophilic lineage within a wider *Metaviridae/*Ty3 retrotransposon clade known as the chromoviruses^81,82^. Furthermore, non-centrophilic relatives of centrophilic *CRM* elements are present in several genomes including those of gymnosperms^82^, mirroring the presence of *ECR-like* elements in *U. pinnatifida*, *D. dichotoma* and *S. ischiensis*. Finally, chromoviruses typically encode a Pol protein that feature a chromodomain fused to the C-terminus of the integrase domain, which presumably targets integration to specific chromatin states^83^. The chromodomain has been replaced by a different putative targeting domain (the “CR” motif) in most *CRM* elements, although a role for this domain in centromeric targeting is yet to be experimentally established. We previously described a C-terminal chromodomain in the pol protein of *ECR-1*^31^. However, both the *ECR* and *ECR*-*like* elements recovered here encode an intact chromodomain, as do other independent non-centrophilic lineages of *Metaviridae*/Ty3 elements in brown algae^31^. Thus, it is unclear if the *ECR* chromodomain is responsible for centromere-targeting, and it is possible that the transition to centrophily evolved following amino acid substitutions in other regions of the integrase domain. Substitutions in the integrase domain between the centrophilic Copia LTR family *Tal1* and the closely related centrophobic family *Evade*, which do not encode chromodomains, was recently shown determine targeting specificity in *Arabidopsis* species^84^.

Similarly, it is also currently unclear how centromere-targeting in the centrophilic Copia elements of *D. dichotoma* and *S. ischiensis* evolved. Phylogenetic analyses suggests that centrophily likely evolved independently in each species, since multiple related families with genome-wide distributions are extant in *S. ischiensis* and brown algae. Alternatively, we cannot rule out that the centrophilic Copia elements represent an ancestral state, with multiple reversions to genome-wide integration patterns occurring among related Copia families. Additional high-quality genomes from non-BACR species will be required to address this question.

Overall, our results capture several themes that are emerging from studies of centromere and retrotransposon co-evolution, including transitions in the identity of the major centrophilic families between species^52,84^. The evolution and ancient association of a chromodomain-encoding lineage of centrophilic *Metaviridae*/Ty3 elements represents a striking case of evolutionary convergence with the *CRM* elements of angiosperms. Spanning at least 160 million years, the ancient centrophily of *ECR* elements may only be exceeded in age by that of the *Bryco* Copia elements of moss centromeres^55^.

Finally, although some centromeres have been lost following specific fusion events (especially in the Ectocarpales), we found that centromere locations were generally conserved among brown algae. This is true even in *D. dichotoma*, suggesting that the transition between Copia and *Metaviridae*/Ty3 centrophilic retrotransposons did not alter centromere locations. Centromere conservation has received considerably less attention in ancestral genome reconstructions, although in some lineages this is impossible due to the presence of holocentric chromosomes. Centromere repositioning refers to the phenomenon of *de novo* centromere evolution without chromosomal rearrangement, which appears to occur relatively frequently in several mammalian and plant lineages^85–88^. Centromere locations have been conserved in the context of the Muller elements for more than 50 MY in *Drosophilia*, although *de novo* centromere evolution was recently shown to have occurred in the *ananassae* subgroup^89^. Similarly, ancestral Brassicaceae centromeres correspond to the extant centromeres in some species such as *Arabidopsis lyrata* and *Capsella rubella*^74^, although some have been lost following fusions in species including *Arabidopsis thaliana*, and other brassica lineages are associated with extensive centromere repositioning^87^. However, few examples of centromere locations that have remained conserved for more than 200 million years have been reported, reinforcing the exceptional nature of macrosynteny in brown algal genomes.

## Methods

### Brown algae culture

Algae materials were cultured in autoclaved natural seawater (NSW) enriched with half-strength Provasoli nutrient solution (Provasoli-enriched seawater; PES) as previously described^30,90^. *C. linearis* (Clin) was grown at 14 °C with the light intensity of 25 μmol photons m^−2^ s^−1^ (8h light/16h dark); *U. pinnatifida* (Upin) male and female strains were grown at 14 °C with the light intensity of 25 μmol photons m^−2^ s^−1^ (12h light/12h dark); *D. herbacea* (Dher) male, female and *D. dudresnayi* (Ddud) strains were grown at 14 °C with the light intensity of 25 μmol photons m^−2^ s^−1^ (12h light/12h dark); *D. dichotoma* (Ddic) male and female strains were grown at 20 °C with the light intensity of 10 μmol photons m^−2^ s^−1^ (16h light/8h dark); *S. ischiensis* (Sisc) were grown at 20 °C with the light intensity of 25 μmol photons m^−2^ s^−1^ (16h light/8h dark). The medium was changed every two weeks.

### Hi-C library preparation

An *in situ* Hi-C protocol of plants was optimized for brown algae^31,61,91^. All strains of *C. linearis*, *U. pinnatifida*, *D. herbacea*, *D. dudresnayi*, *D. dichotoma*, *S. ischiensis* were cultivated in controlled lab conditions^25^ and material was collected using a 40 µm filter and fixed in 2 % (vol/vol) formaldehyde for 30 min at room temperature, and the cross-linking reaction was quenched with 400 mM glycine. Approximately 50 mg of frozen algal tissue was ground in liquid nitrogen using a pre-chilled mortar and pestle. The resulting fine powder was suspended in 5 ml nuclei isolation buffer (0.1% triton X-100, 125 mM sorbitol, 20 mM potassium citrate, 30 mM MgCl_2_, 5 mM EDTA, 5 mM 2-mercaptoethanol, 55 mM HEPES at pH 7.5) with 1X protease inhibitor, and transfer into a 7 mL Tenbroeck potter. Grind 10 times slowly on ice; then transfer the solution into in several 2 ml VK05 tube, homogenized by Precellys Evolution beads homogenizer (Bertin technologies, 7800 rpm, 30s each time, 20s pause each grinding cycle, repeat 5 times). Over 1 million nuclei were isolated and digested overnight by Dpn II, DNA ends were labeled with biotin-11-dCTP (Jena Bioscience, cat. no. NU-809-BIOX-L) at 22°C for 4h in thermomixer (shake at 900 rpm, alternating 30 seconds on and 4 minutes off), then ligated by T4 DNA ligase (Thermo Scientific, cat. no. EL0012) at 22°C for 4h. The purified Hi-C DNA was sheared for 60 seconds by covaries E220 evolution (Peak incident power:175W, duty cycle: 10, intensity: 190, cycles per burst: 200) and libraries were prepared using the NEBNext Ultra II DNA Library Prep Kit (NEB, cat. no. E7645) following the standard protocol, DNA concentration was quantified using a Qubit™ 4 Fluorometer (Invitrogen), and the average size of the library was detected by Bioanalyzer (Agilent Technologies), the final library was sequenced with 150 bp paired-end reads on an Illumina HiSeq 2000 platform at MPI and Novaseq X at Azenta. To check library quality and adjust sequence depth, an aliquot of library was sent for test sequencing. Two biological replicates were performed for each strain.

### High-Molecular-Weight (HMW) genomic DNA extraction and Nanopore sequencing

HMW gDNA was extracted from *U. pinnatifida* strains, *D. dichotoma* female strains and *D. dudresnyi* strains using the NucleoBond® HMW DNA kit (Macherey-Nagel, cat. no. 740160.20) in combination with Lysis Buffer CF (Macherey-Nagel, cat. no. 740946). Approximately 300 mg of frozen algal tissue was ground in liquid nitrogen using a pre-chilled mortar and pestle. The tissue was grinded into fine powder and suspended in Lysis Buffer CF, gently inverted to ensure homogenous mixing, followed by incubation at 37 °C for 10 minutes. Subsequently, the following reagents were added sequentially, with gentle inversion after each addition: 5 mL of 5 M NaCl, 200 µL of 0.5 M EDTA, 4 mL of 10% CTAB in 0.7 M NaCl, 200 µL of 10% Triton X-100, 400 µL of Proteinase K (20 mg/mL), and 200 µL of RNase A. The extraction process then followed the manufacturer’s protocol for HMW DNA isolation. DNA concentration was quantified using a Qubit™ 4 Fluorometer (Invitrogen), and fragment size distribution was assessed using the FEMTO Pulse system (Agilent Technologies) to confirm the integrity of HMW gDNA. For long reads sequencing, 1 µg of HMW gDNA was used to prepare libraries following the standard protocol of the Oxford Nanopore Technologies (ONT) ligation sequencing kit SQK-LSK110 (*D. dichotoma, D. dudresnayi*) and SQK-LSK114 (*U. pinnatifida*). Sequencing was performed on MinION R9.4 (*D. dichotoma* and *D. dudresnayi*) and PromethION2 R10.4 (*U. pinnatifida*) platforms. Base- calling was performed by ONT dorado v0.3.4 (https://github.com/nanoporetech/dorado).

### *De novo* genome assembly and scaffolding

ONT long reads draft assemblies were performed by canu^92^, flye^93^, NextDenovo^94^ and Shasta^95^ long read assemblers with default settings. The best draft assembly for each species was chosen based on the contig fragmentation level, N50, repeated sequences behavior, the re-mapped read coverage and the overall assembly length. We improved nanopore-based draft genome assemblies to chromosome-level resolution using Hi-C data. Specifically, *C. linearis*, and *S. ischiensis* draft assemblies were scaffolded using the 3D-DNA pipeline,^96^ while the *D. dichotoma* female assembly was improved using the HapHiC pipeline (v1.0.7)^97^. Mapping in situ Hi-C data to existing *D. herbacea* male and female genomes, as well as to *D. dichotoma*, revealed multiple mis-assemblies in the reference scaffolds. To address these, we applied the 3D-DNA pipeline to the draft assemblies of *D. herbacea* male, female and *D. dudresnayi*^26^ and the HapHiC pipeline (v1.0.7) to the *D. dichotoma* male genome^26^ using Hi-C datasets generated in this study. Mis-assemblies and chromosome rearrangements were corrected manually using Juicebox^45^. Overall, we obtained near telomere-to-telomere (T-to-T), chromosome-level assemblies for all species included in this study. In the updated assemblies, chromosomes were renamed and artificially oriented according to previous reference versions. Genome completeness was assessed using BUSCO (v5.8.2) with the eukaryota_odb10 dataset^38^.

### Genome annotation and macrosynteny analysis

Gene annotations were lifted over from the previous gene models to the new assembled genomes used Liftoff (v1.6.1)^98^, and repeat was re-annotated by Earl Grey^99^ and manually curated with *Ectocarpus* sp7 TE library^40^. Homologous syntenic blocks were built with GENESPACE^41^ R package and visualized by LINKVIEW2^100^.

### Hi-C data processing

Hi-C data was processed by Juicer^101^ pipeline with default parameters, Hi-C reads were mapped to chromosome-level assemblies of each species with BWA, after alignment, chimera handing, merge, sort and removed duplicate reads pairs, juicer format contact maps(.hic) were generated with resolutions of 5kb, 10kb, 25kb, 50kb, 100kb and 250kb and visually inspected in Juicebox^45^. HiCRep^42^ was used to assess the Hi-C data reproducibility between replicates with stratum-adjusted correlation coefficient (SCC) method at resolutions of 5, 10, 25, 50, 100 and 250kb. The SCC scores were averaged across chromosomes. Biological replicates were merged to get a higher resolution Hi-C matrix. Juicer format pair files ‘merged_nodups.txt’ was convert to pairs with ‘merged_nodup2pairs.pl’. The distance law represents decay of the average contact frequency was calculated directly from pair files with HiContacts^43^ R package, the most informative genomic distance from 10 kb to 1 Mb was plot for each sample separately.

### Compartment A/B

Chromatin compartments were inferred from based on the plaid interaction patterns in Hi-C contact matrices, which reflect spatial segregation of A and B compartment (open and close chromatin). To annotate compartment A/B, juicer format matrix‘.hic’ files with a mapping quality score (MAPQ) ≥ 30 were converted to .cool format by hic2cool (https://github.com/4dn-dcic/hic2cool, v1.0.1) and matrix balancing was performed using the cooler (v1.0.1) ^102^ by iterative correction and eigenvector (ICE) method. Eigenvector decomposition was conducted using the cooltools^103^ (v0.4.0) eigs-trans function to calculate the first principal component (E1) at multiple resolutions. The sign of the E1 values was corrected based on their correlation with genomic features: regions with higher GC or gene density were designated as compartment A (positive E1), while regions with lower density were assigned to compartment B (negative E1). To refine compartment annotations, E1 signal tracks were visually inspected alongside Hi-C contact maps with KR normalization^104^ in Juicebox^45^. Manual validation focused on ensuring concordance between compartment signals and local intra- and inter-chromosomal interaction patterns. Saddle plots were generated following the instruction (https://cooltools.readthedocs.io/en/latest/notebooks/compartments_and_saddles.html), 95% of the genome was divided into 38 groups based on digitized eigenvector values, and we calculated the average observed/expected contact frequencies in each pair of group^103^. From each saddle plot, we extracted the average interaction frequencies within B–B compartments (top left) and A–A compartments (bottom right). Compartment strength was quantified as the ratio of (AA + BB) / (AB + BA)^103^.

### Insulation score and boundary calculation

Insulation score reflects the aggregation level of contact frequency map, high insulation is classified as chromatin boundaries in plant and animal^105^. To annotate insulation, cooltools^103^ (v0.4.0) was used with balanced cooler format matrix‘.mcool’ files as input. Using Hi-C contacts matrix with a mapping quality score (MAPQ) ≥ 30 at 10 kb resolution, we tested multiple window sizes (50, 100, 200, 300, 400, and 500 kb). Based on visual inspection with Hi-C matrix in Juicebox^45^, the 100 kb window size was selected for downstream analysis. Strong and weak boundaries were identified using the parameters ‘--threshold Li --min-dist-bad-bin 2’.

### Identification of centromeric retrotransposons

Each genome was first queried via tblastn searches^106^ using the ECR-1 pol protein from the *Ectocarpus* sp. 7 centrophilic retrotransposon^31^. The hits in each genome were then manually curated and consensus sequences were produced for each corresponding family following the method of Goubert et al. ^107^. Briefly, the nucleotide sequences were extracted with flanking sequence, clustered based on similarity, and then reduced to a single full-length consensus sequence based on alignments that extended from the start of the left long terminal repeat to the end of the right long terminal repeat.

As a second approach, broad centromeric regions of ∼1 Mb were visually defined from the Hi-C contact maps for each species (see Fig. 4). As performed previously for *Ectocarpus* sp. 7 ^31^, putative centromeric repeats from the automated Earl Grey repeat libraries were defined based on i) their enrichment in the centromeric regions relative to non-centromeric regions, and ii) their presence in multiple, if not all, centromeric regions. Any other repeat families beyond the *ECR* families that met these criteria were also manually curated following the same process, which included the centrophilic Copia families from *D. dictyota* and *S. ischiensis*.

If present, redundant repeat models from the automated Earl Grey libraries were replaced with the manually curated models, and the updated libraries were then used to annotate repeats against the relevant genome using RepeatMasker v4.1.6 (repeatmasker.org). The *Ectocarpus* sp. 7 genome was instead masked using the available curated library^40^. Additional satellite repeats were identified in each genome using Tandem Repeats Finder v4.09.1 and the parameters “2 7 7 80 10 50 2000”, followed by extracting repeats with monomer lengths > 10 bp. If TE and satellite repeat annotations overlapped, TEs were given precedence. The genomic densities of TEs (including centrophilic families and related *ECR-like* families) and satellites were then calculated in nonoverlapping sliding windows.

For *ECR* and *ECR-like* families, gag and pol protein sequences were manually extracted from the nucleotide consensus sequences and aligned using mafft v7.525 and the “L-INS-i” model^108^. Alignment gaps were filtered out using trimAl v1.4rev22 and the model “gappyout”^109^. A maximum likelihood phylogeny was then produced using IQ-TREE v2.3.0 with ultrafast bootstrapping and ModelFinder (“-bb 1000 -m MFP”)^110^. For Copia, Gag and Pol protein sequences were extracted from the consensus sequences for the centrophilic families and for all curated *Ectocarpus* sp. 7 families^40^. The *D. dichotoma* reverse transcriptase domain was used to perform tblastn searches against the automated Earl Grey repeat models for each species, corresponding protein sequences were extracted and a preliminary phylogeny was produced using the method above. This phylogeny was used to identify a candidate set of Copia families that are closely related to the centrophilic families, and full-length gag and pol sequences were extracted for each of these families (with manual curation performed if required to extend consensus sequences or correct for frame-shifts). A final phylogeny using the resulting gag and pol protein sequences was then produced using the same method as for *ECR*/*ECR-like* elements, with more distantly related Copia families from the curated *Ectocarpus* sp. 7 library forming the outgroup.

### Ancestral genome reconstruction

Ancestral genome reconstruction was performed using AGORA^60^ and the following genomes and annotations: *Ectocarpus* sp. 7^31^, *C. linearis*, *U. pinnatifida*, *D. herbacea*, *D. dictyota* (all this study), and *S. promiscuus* and *F. serratus* ^25^. Single copy orthologs present in all genomes (n=4,847) were defined using OrthoFinder v2.5.5 (“-S diamond_ultra_sens”)^111^. Contiguous Ancestral Regions (CARs) flanking the centromeres were then extracted and the genes present in these CARs were mapped onto the chromosomes (see Figure 5).

Rearrangements were manually curated using the GENESPACE macrosynteny analysis^41^. A second OrthoFinder run including *S. ischiensis* proteins was performed, and orthologs between *D. herbacea* and *S. ischiensis* were extracted. One-to-many and many-to-many orthology relationships were retained if all of the genes were on the same chromosome in both species. The *S. ischiensis* genes were then colored according to the chromosome of their ortholog(s) in *D. herbacea*.

## Acknowledgements

We thank Remy Luthringer, Masakazu Hoshino, Kenny A Bogaert, Andrea Belkacemi, Dorothe Koch and Anagha Kerur, for assistance with algal cultures; Jaruwatana Sodai Lotharukpong and Romy Petroll for help with phylogenetic trees; Jeromine Vigneau, Michael Borg, Erica Dinatale and Zhigui Bao for discussions.

## Funding

This research was funded by the Max Planck Society, European Research Council, grant 864038 (SMC), the Gordon and Betty Moore Foundation (SMC). RC was recipient of a Marie Skłodowska-Curie Postdoctoral Fellowship (grant 101109906). Pengfei Liu is thankful to the International Max Planck Research School ‘From Molecules to Organisms.’

## Author contributions

Conceptualization: SMC. Methodology: SMC, RC, PL, CL. Investigation: PL, RL, FBH, RC. Visualization: PL, RC, FBH. Funding acquisition and project administration: SMC. Supervision: SMC, CL, FBH. Writing (original draft): SMC, PL, RC. Writing (review & editing): SMC.

## Competing interests

Authors declare that they have no competing interests.

## Data and materials availability

All data are available in the main text or the supplementary materials.

## Supplemental Figures

**Fig. S1.**
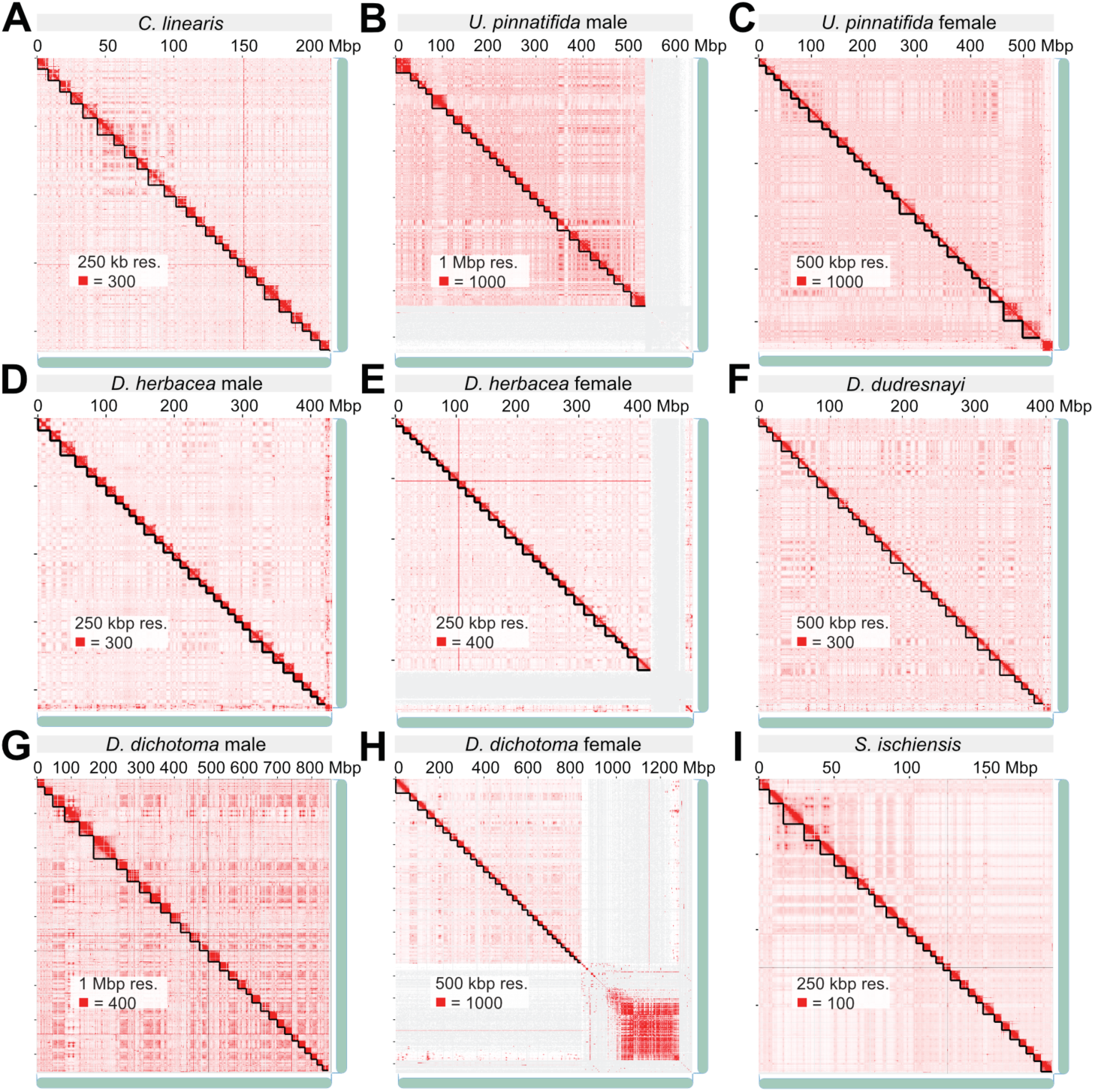
(**A**-**I**) Genome assemblies for each strains by nanopore long reads and Hi-C reads, chromosomal boundaries and prominent interactions of telomere clustering and centromere clustering are clear seen in Hi-C maps for each species in Juicebox^45^.

**Fig. S2.**
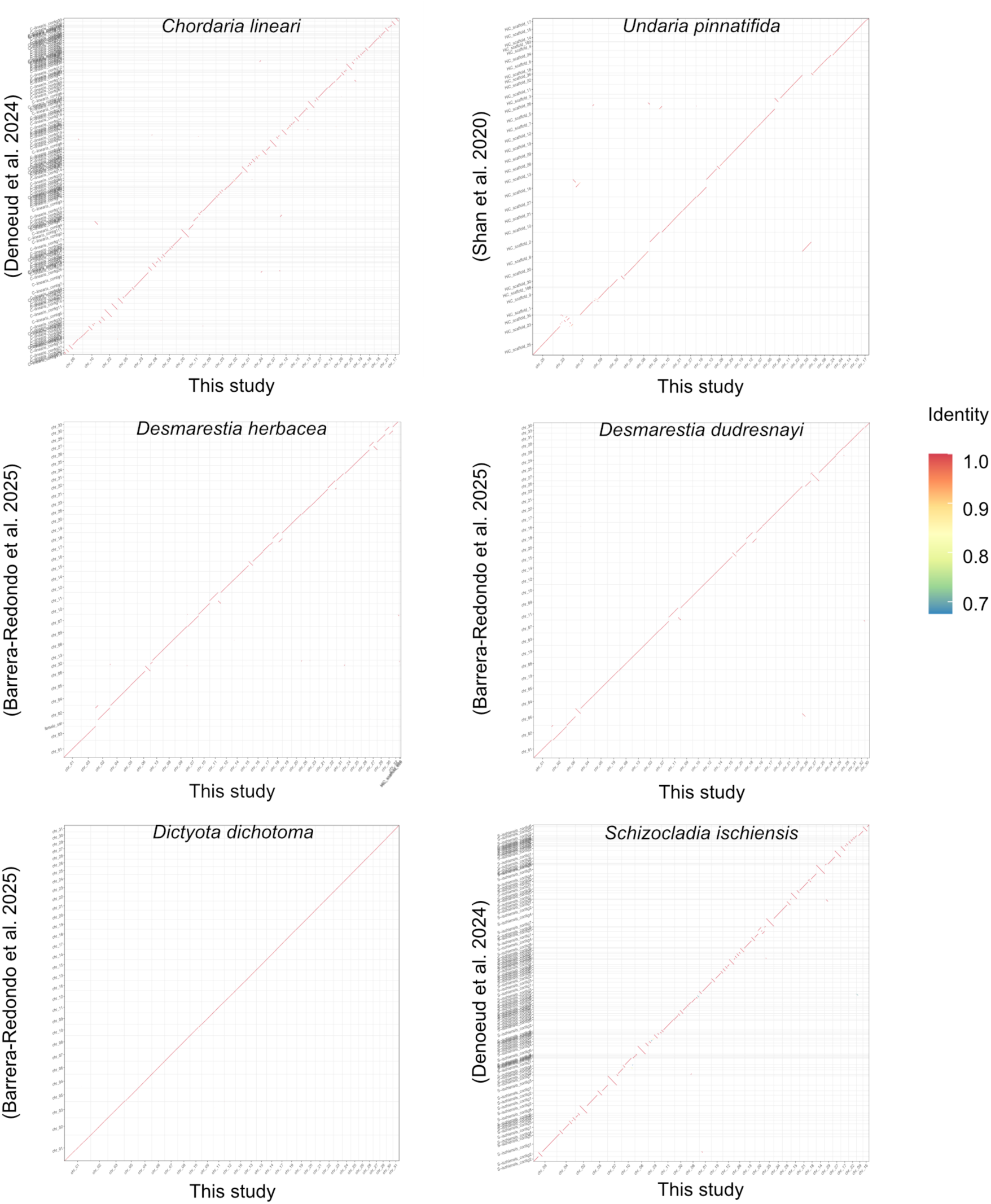
Dotplot comparison of previous and current genome assemblies. Whole-genome pairwise alignments were performed using minimap2^112^, and the resulting alignments were visualized with paf2dotplot (https://github.com/moold/paf2dotplot). The generated dotplots indicate high sequence identity between the old and new assemblies

**Fig. S3.**
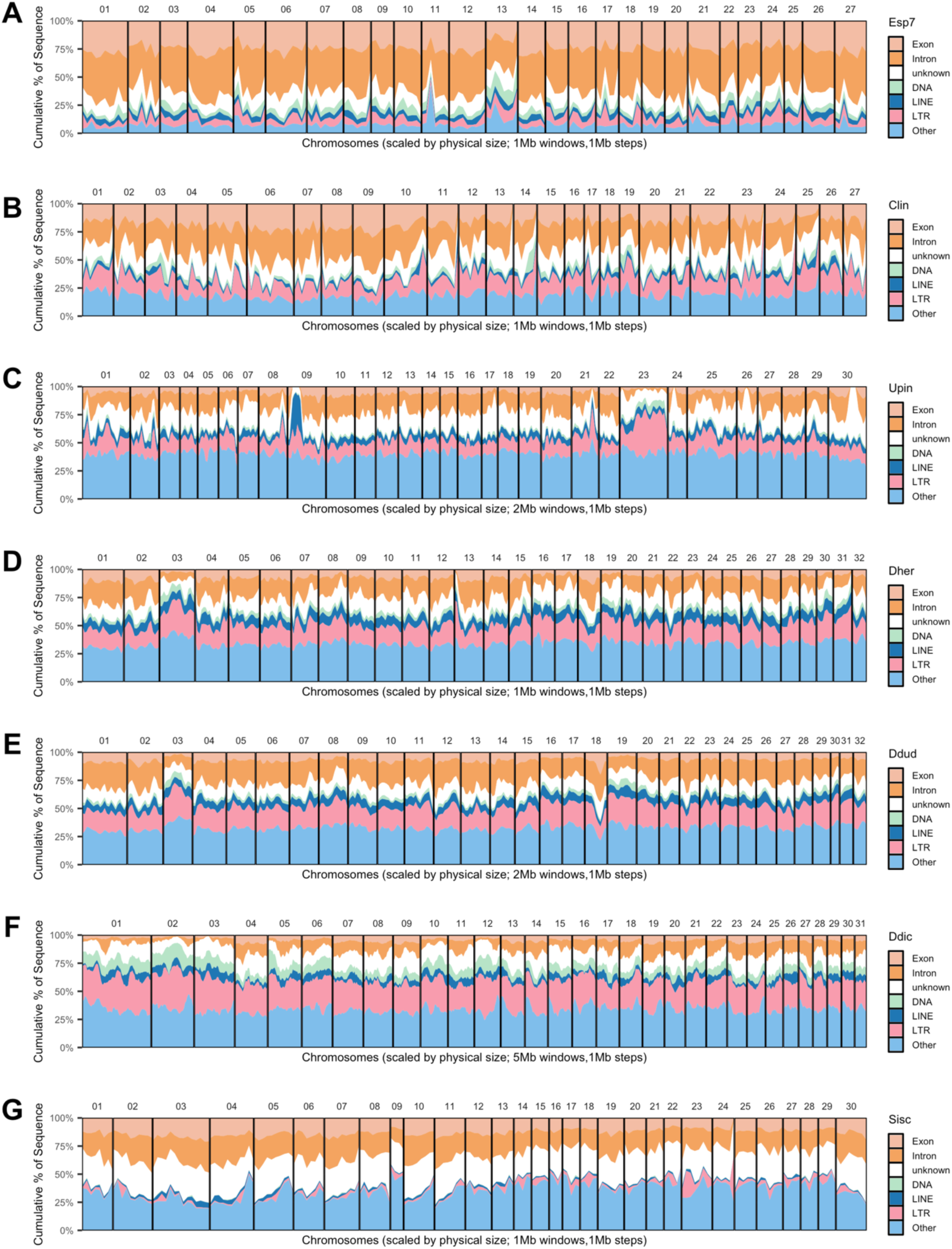
(**A-G**) Landscape of exon, intron, and TE families (DNA, LINE, LTR and others, others include simple repeats, satellite, unclassified DNA, etc) for each species. (Esp7 = *Ectocarpus* sp. 7, Clin = *Chordaria linearis*, Upin = *Undaria pinnatifida*, Dher = *Desmarestia herbacea*, Ddud = *Desmarestia dudresnayi*, Ddic = *Dictyota dichotoma*, Sisc = *Schizocladia ischiensis*).

**Fig. S4.**
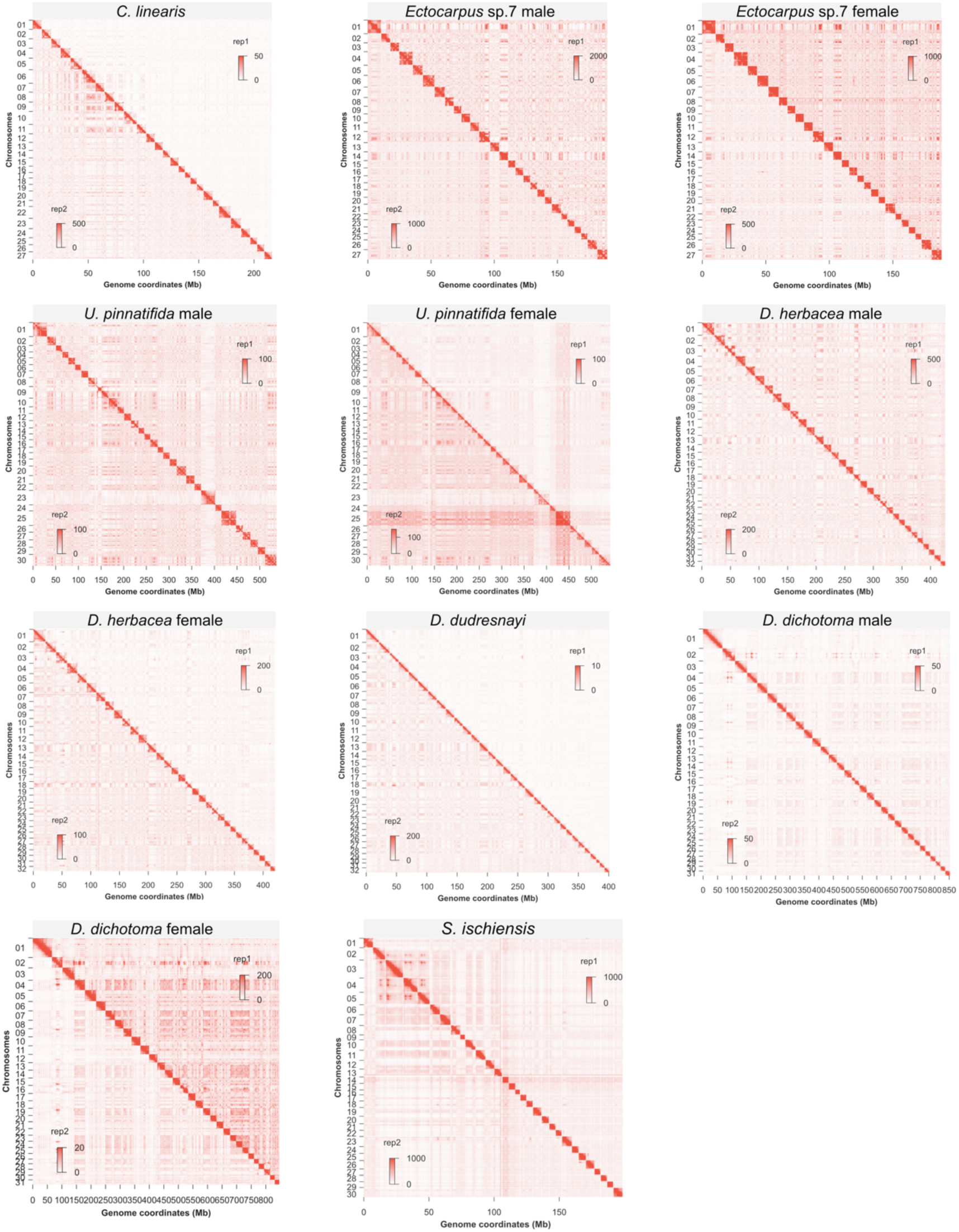
Biological replicates of whole genome-wide Hi-C maps for each species visualized in Juicebox^45^ at 250k resolution

**Fig. S5.**
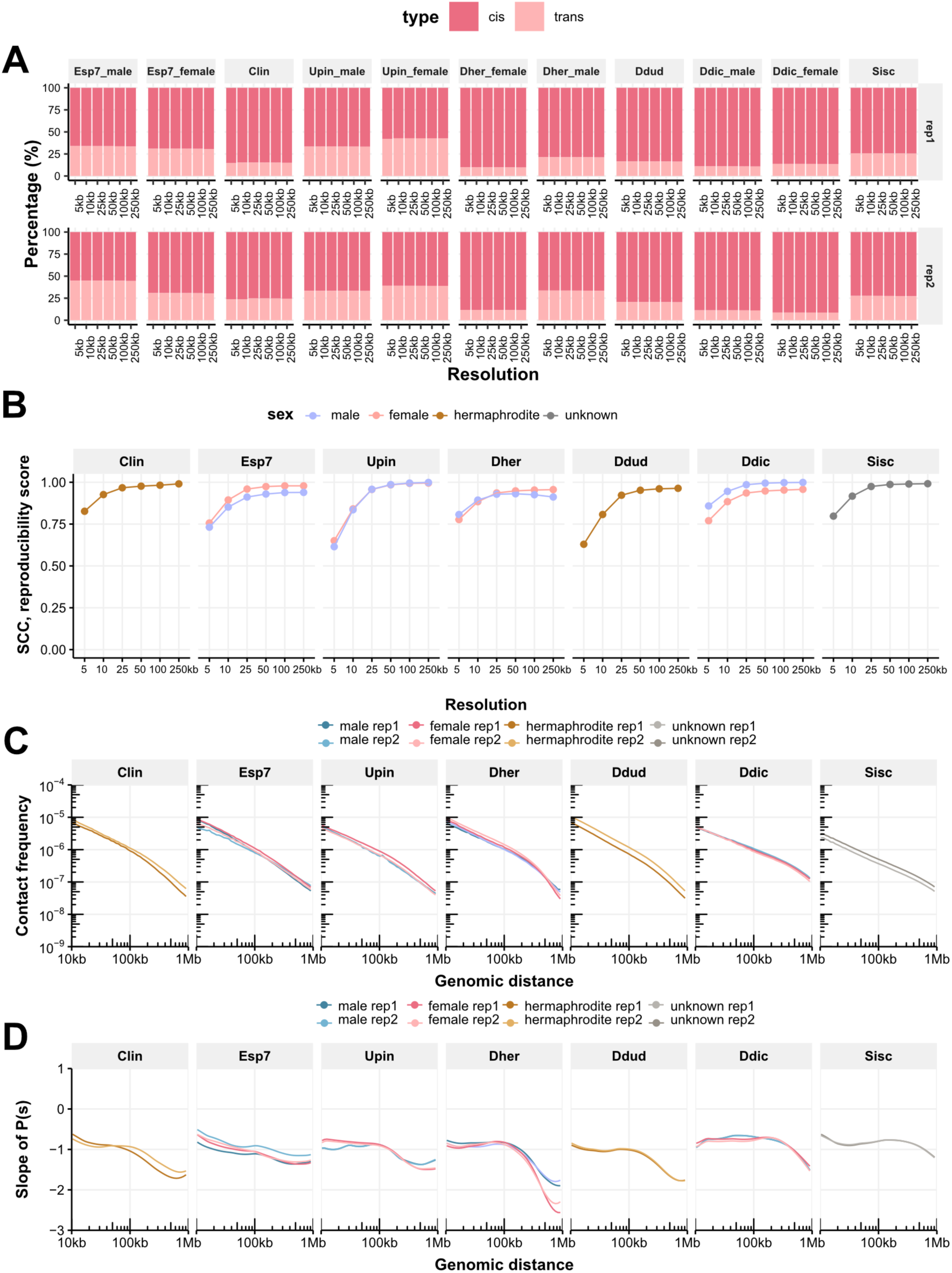
Hi-C data statistics. (**A**) Percentage of cis and trans interactions at different resolutions. (**B**) corrections score between biological replicates, and (**C**) Interaction frequency curve of distance law and (**D**) slope of distance law for each replicate of each species

**Fig. S6.**
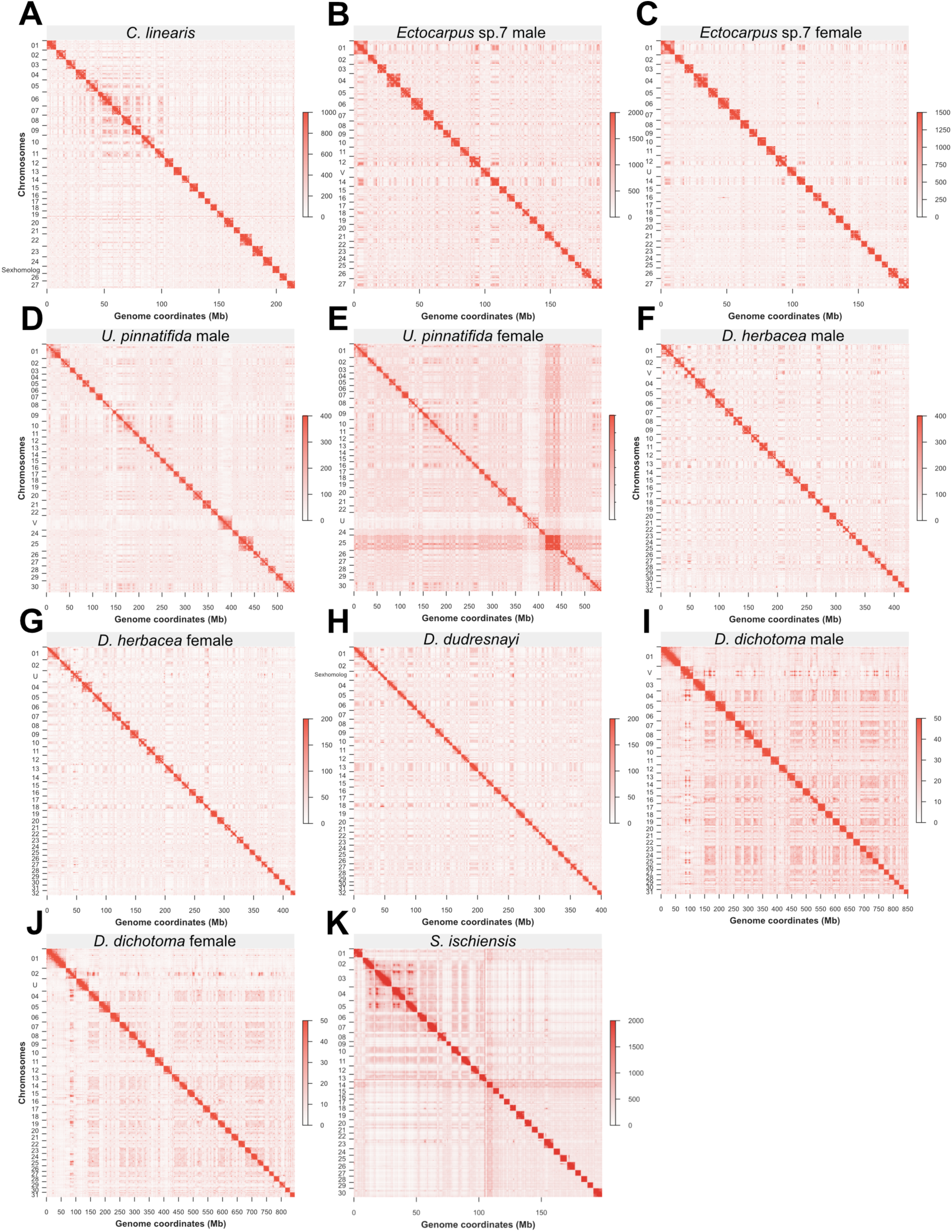
Final whole genome-wide Hi-C maps for each species at 250k resolution with merged Hi-C data.

**Fig. S7.**
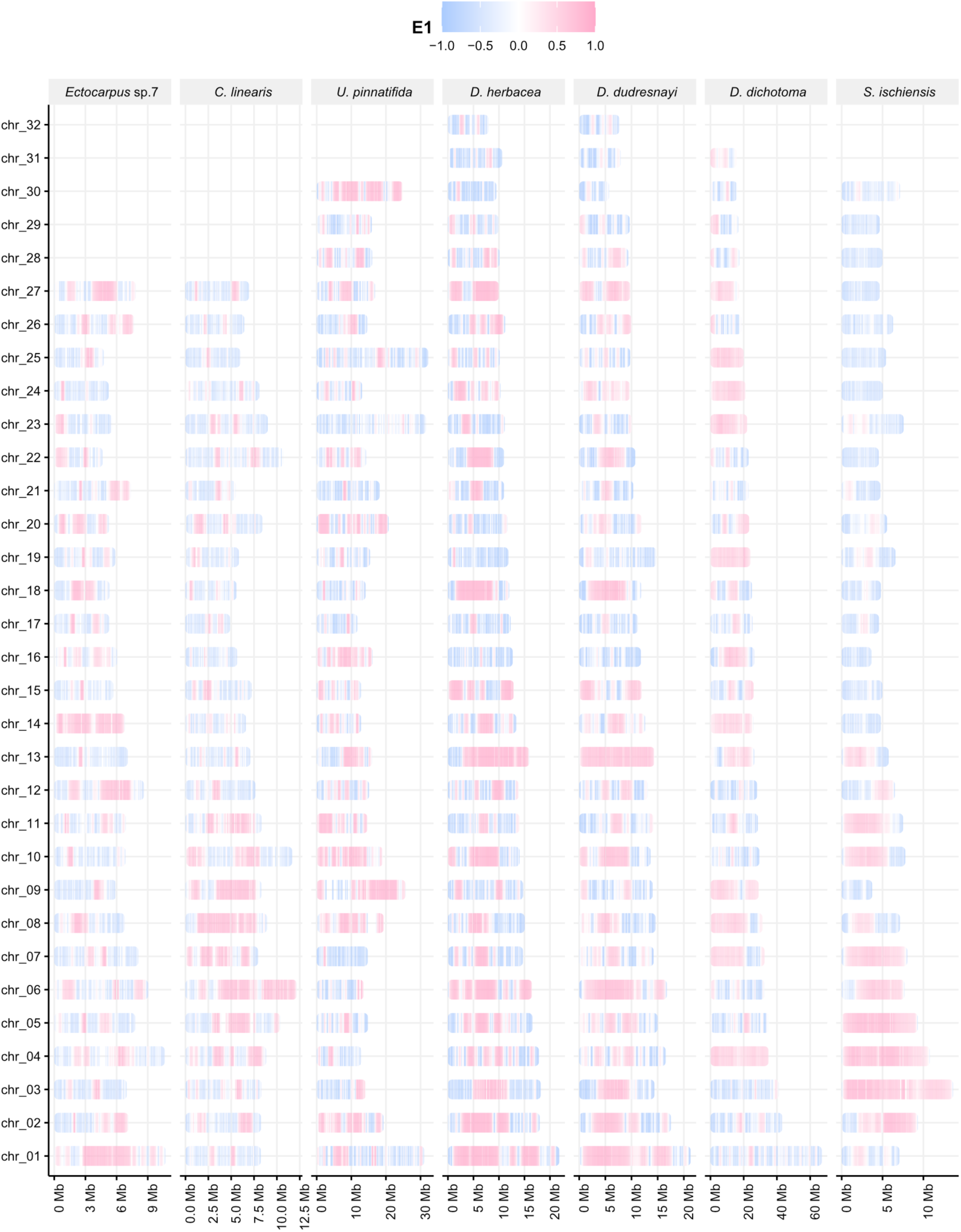
A/B compartment distribution on each chromosome for each species (E1 is the first eigenvector value).

**Fig. S8.**
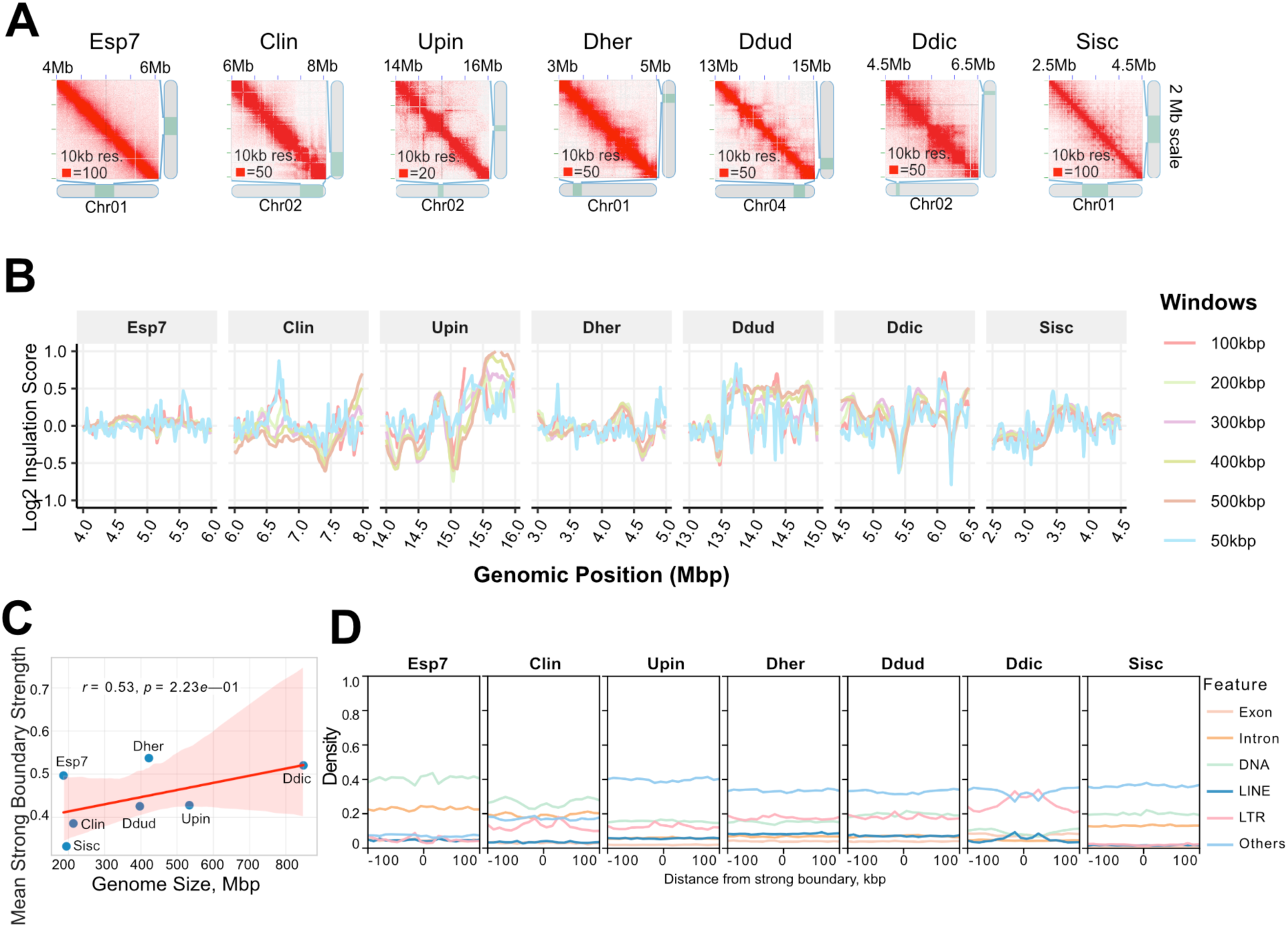
Insulation and boundaries profiles across species. (**A**) A 2 Mb-scale Hi-C matrix for each species. (**B**) Insulation scores were calculated using 10 kbp resolution Hi-C contact matrices with multiple sliding window sizes (50 kbp, 100 kbp, 200 kbp, 300 kbp, 400 kbp, and 500 kbp). To compute insulation profiles, a diamond-shaped window was slid along the genome diagonal, with one corner anchored on the main diagonal of the contact matrix. For each genomic position, the sum of contacts within the window was calculated, with lower scores indicating stronger insulation boundaries. (**C)** Correlation between extracted mean boundary strength score and genome size, the shaded region represents the 95% confidence interval around the regression line. (**D**) Exon, intron and TEs enrichment on boundaries upstream and downstream at 10k window size. Numbers below brackets represent the number of boundaries. The lower and upper hinges of the box correspond to the first and third quartiles (the 25th and 75th percentiles). The upper whisker extends from the hinge to the largest and smallest values no further than 1.5x IQR from the hinge (Inter-Quartile Range, distance between the first and third quartiles). Significant was determinant by two-sample Wilcoxon rank sum test.

**Fig. S9.**
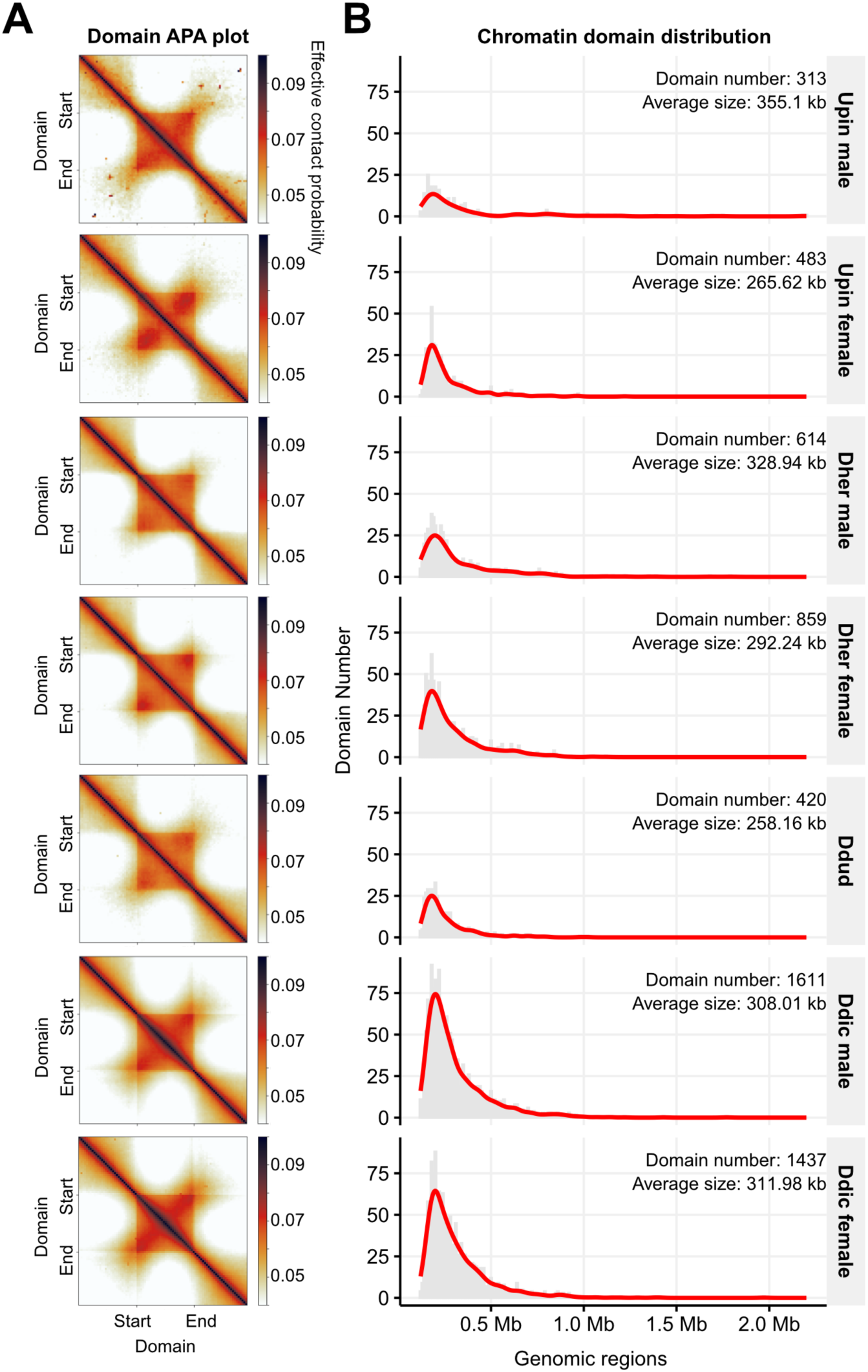
Chromatin domains profile for species. (**A**) Aggregate chromatin domain plot of normalized Hi-C matrix at 10kb resolution by FAN-C^113^. For each TAD, the contact map from (start - length) to (end + length) was rescaled to a 90×90 matrix and averaged. The TAD body corresponds to the central pixels. (**B**) Domain size distribution in each species.

**Fig. S10.**
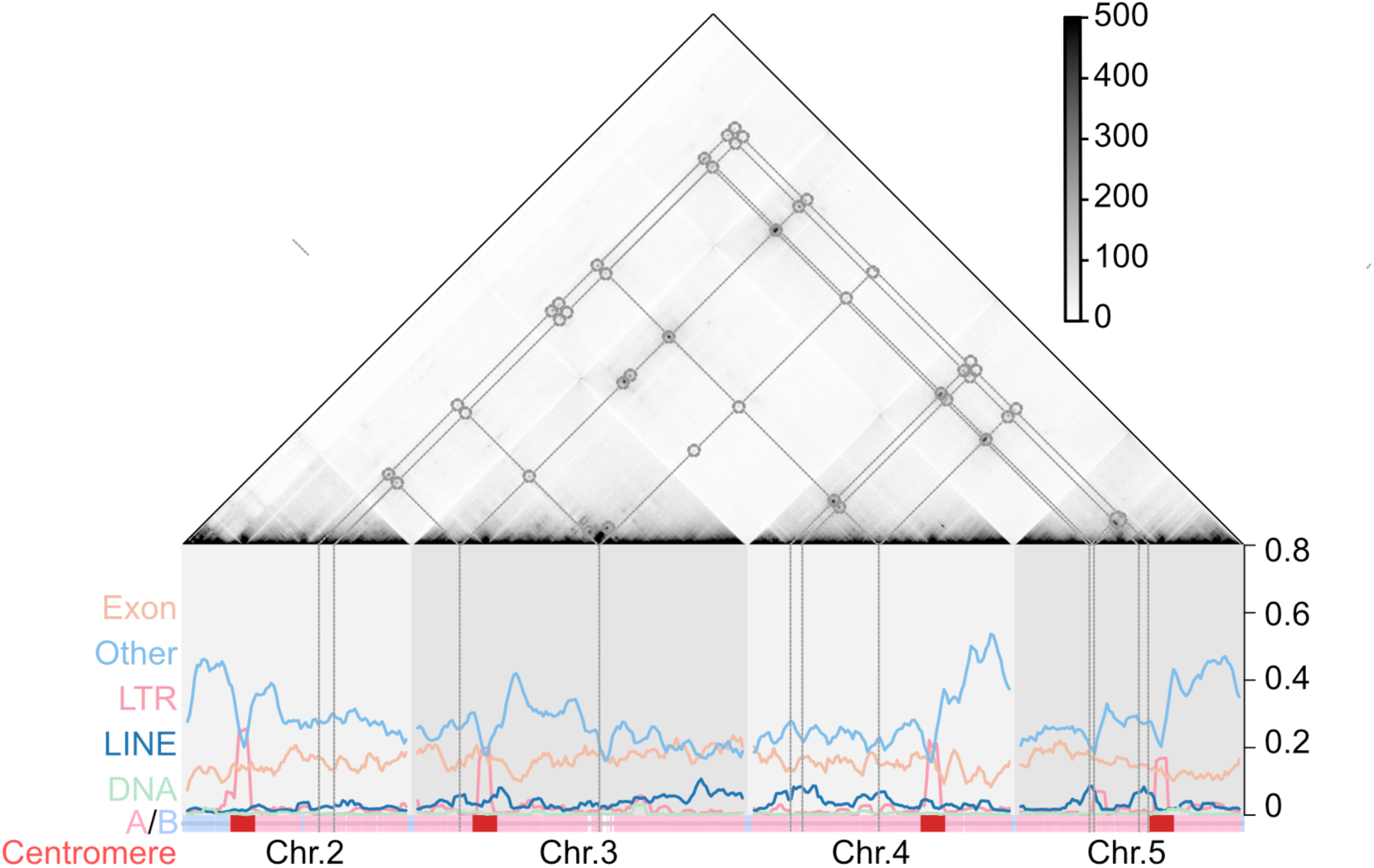
Example shows prominent inter-chromosomal interactions of a sub Hi-C matrix of *S. ischiensis* at 25k resolution, profiled with fraction of TEs family (LINE, DNA, LTR and others), exon in 500kb window size, ideogram shows compartment A/B and centromere annotations.

**Fig. S11.**
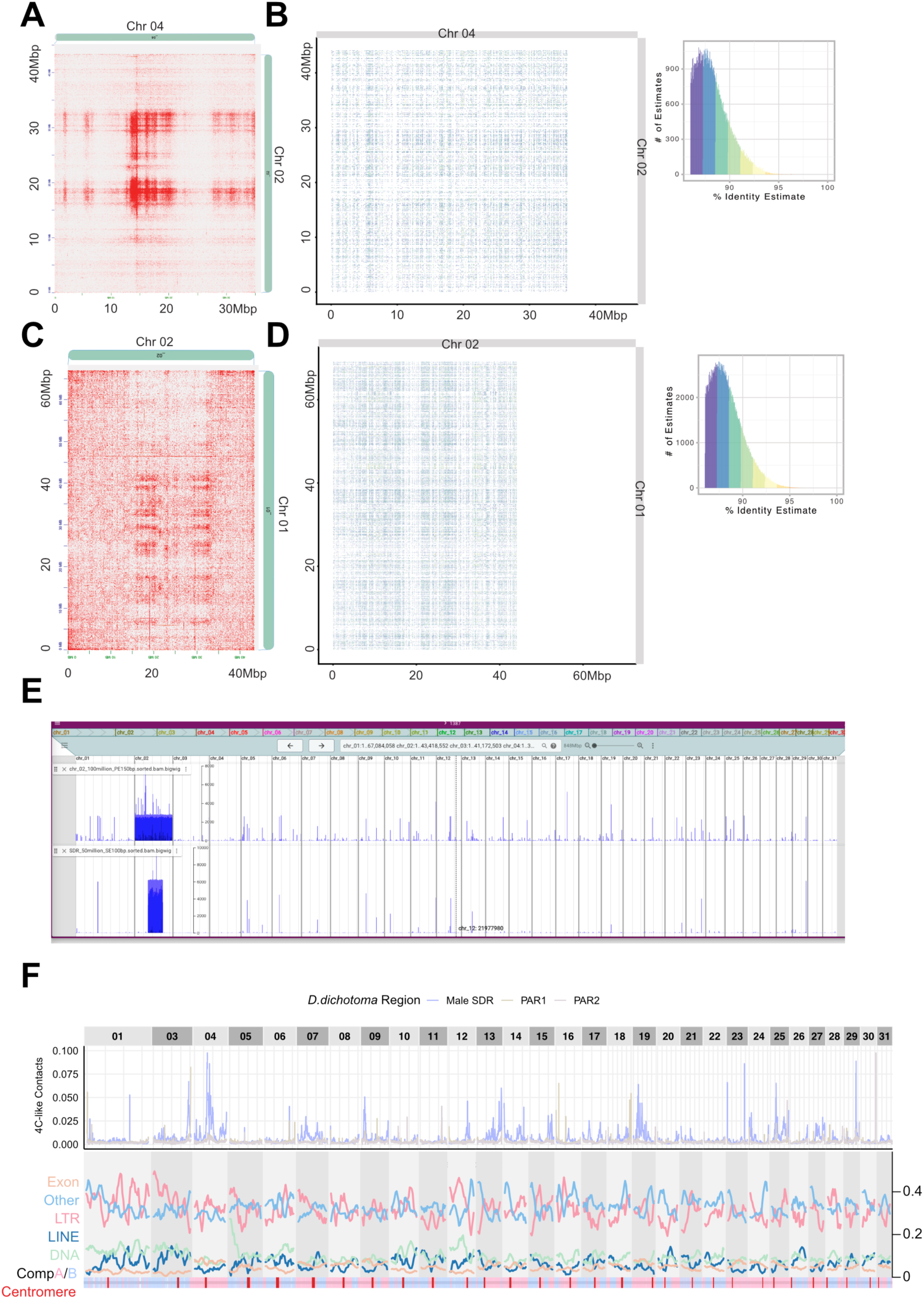
Validation of long-range interactions between sex chromosomes (SDRs) and autosomes are not mapping artifacts and interaction patterns are not affected by TEs. (**A**) and (**C**) Examples showing strong inter-chromosomal interactions involving the SDR on *D. dichotoma* male genome. (**B**) and (**D**) corresponding self-identify plots reveal low sequence similarity between the sex chromosomes and autosomes by ModDotPlot^114^. (**E**) Coverage of uniquely mapped reads: simulated 100 million paired-end (PE) 150 bp reads of the entire sex chromosome (upper), and simulated 50 million single-end (SE) 100 bp reads of male SDR (lower), shown on *D. dichotoma* male genome. (**F**) Genome-wide virtual 4C profile of contacts between PAR1, SDR, and PAR2 of sex and autosomes at 100kb Hi-C resolution and corresponded each TEs family density, compartment A/B and centromere (in red) profiles for *D. dichotoma* at 5Mb window size.

**Fig. S12.**
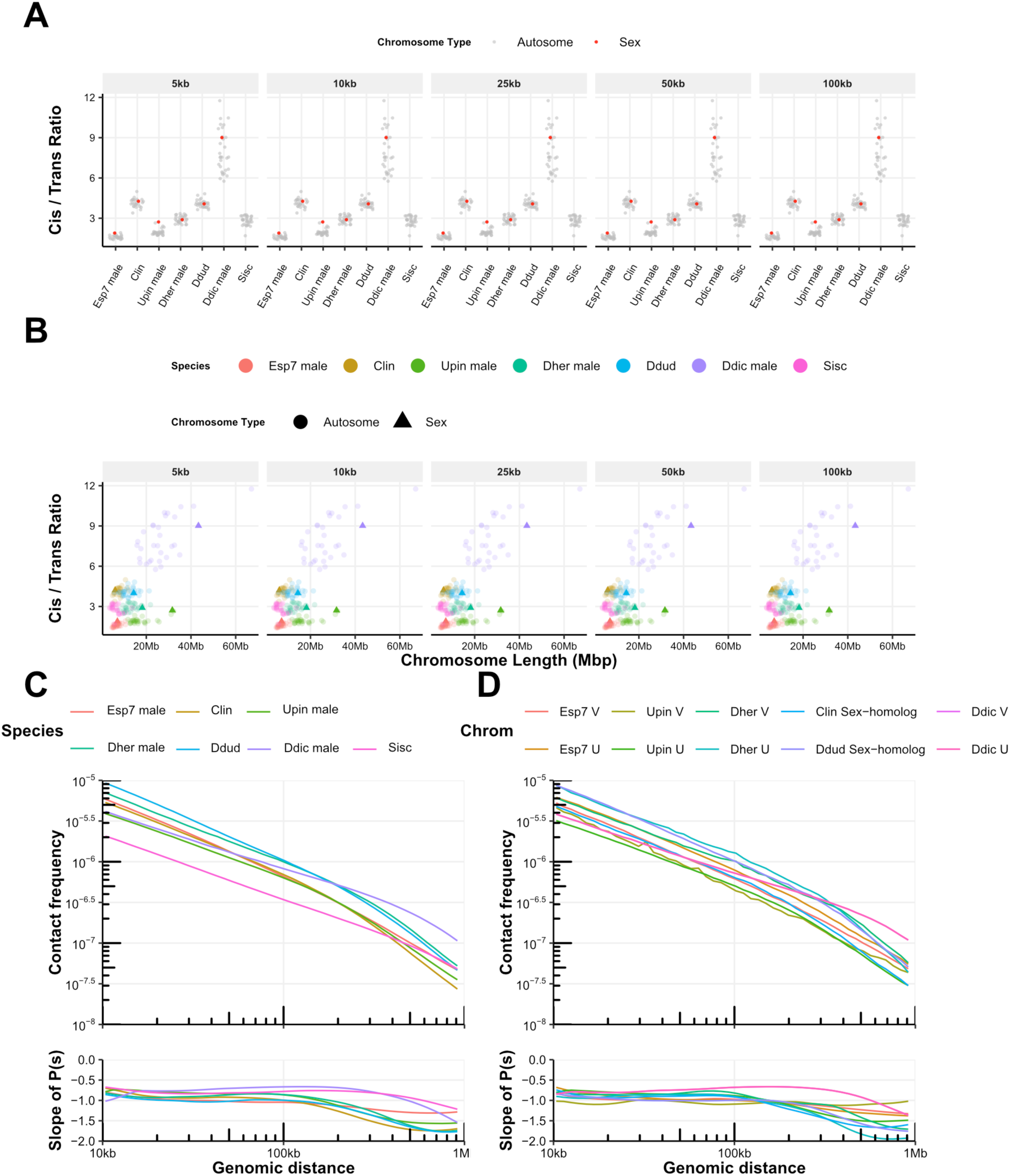
Trans/cis interaction frequency ratio for (A) each chromosome and (**B**) as a function of chromosome size, shown for each species at resolutions of 5 kb, 10 kb, 25 kb, 50 kb, 100 kb, and 250 kb. (**C**) Distance-dependent interaction frequency (Ps) and corresponding slope for each species) and (**D**) sex chromosomes (Chr V and Chr U) and sex-homologs.

**Fig. S13.**
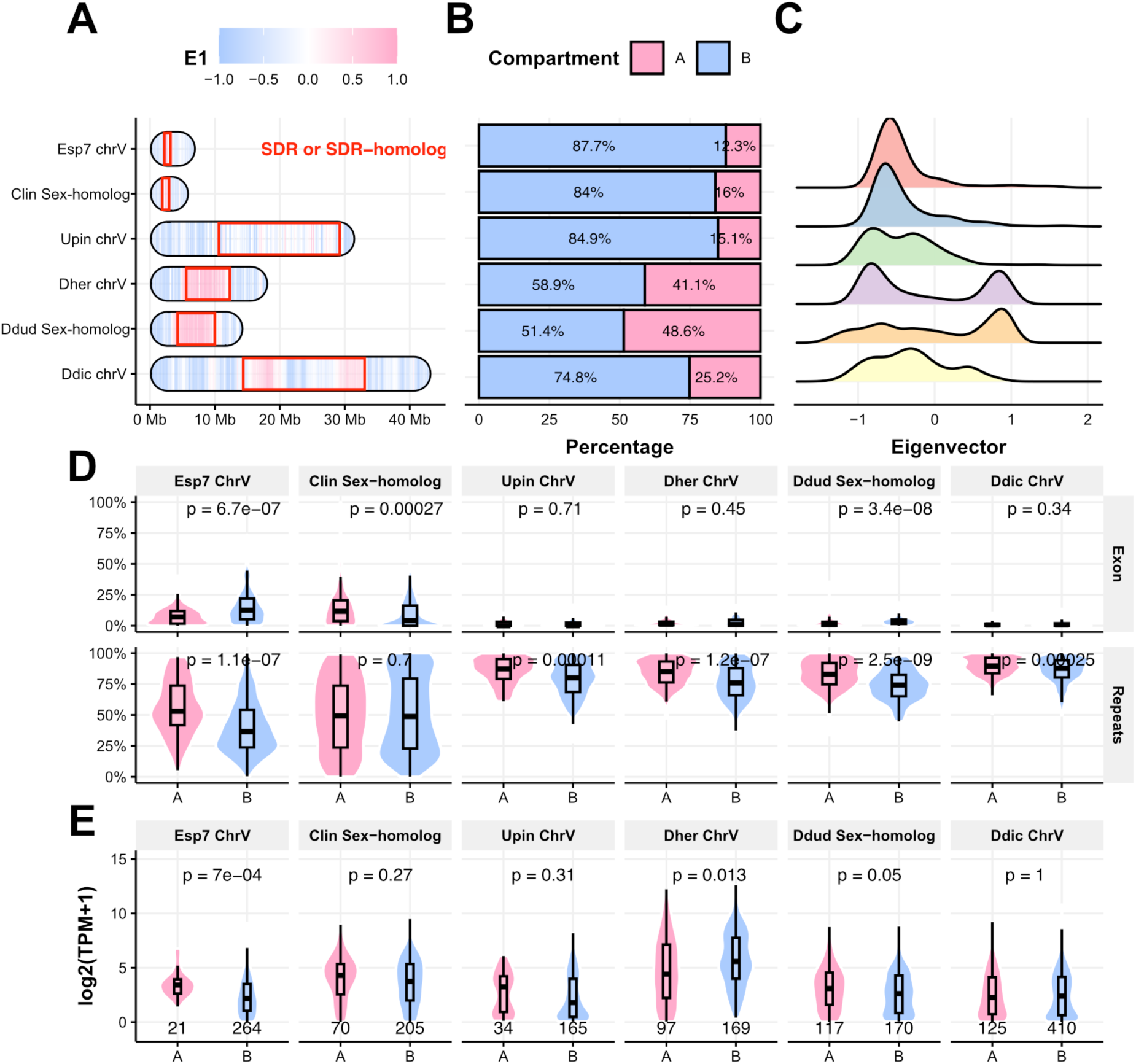
Compartmentalization profile of sex chromosome (Chr V and Chr U) and sex homologs. (**A**) ideogram plot for eigenvector profiles, (**B**) percentage of A or B compartment, (**C**) eigenvector values distribution (**D**) exon and repeat density and (**E**) gene expression in A / B compartment for sex chromosomes and sex homologs of each species. Numbers below brackets represent the number of genes. The lower and upper hinges of the box correspond to the first and third quartiles (the 25th and 75th percentiles). The upper whisker extends from the hinge to the largest and smallest values no further than 1.5x IQR from the hinge (Inter-Quartile Range, distance between the first and third quartiles). Significance was determined by a two-sample Wilcoxon rank sum test.

**Fig. S14.**
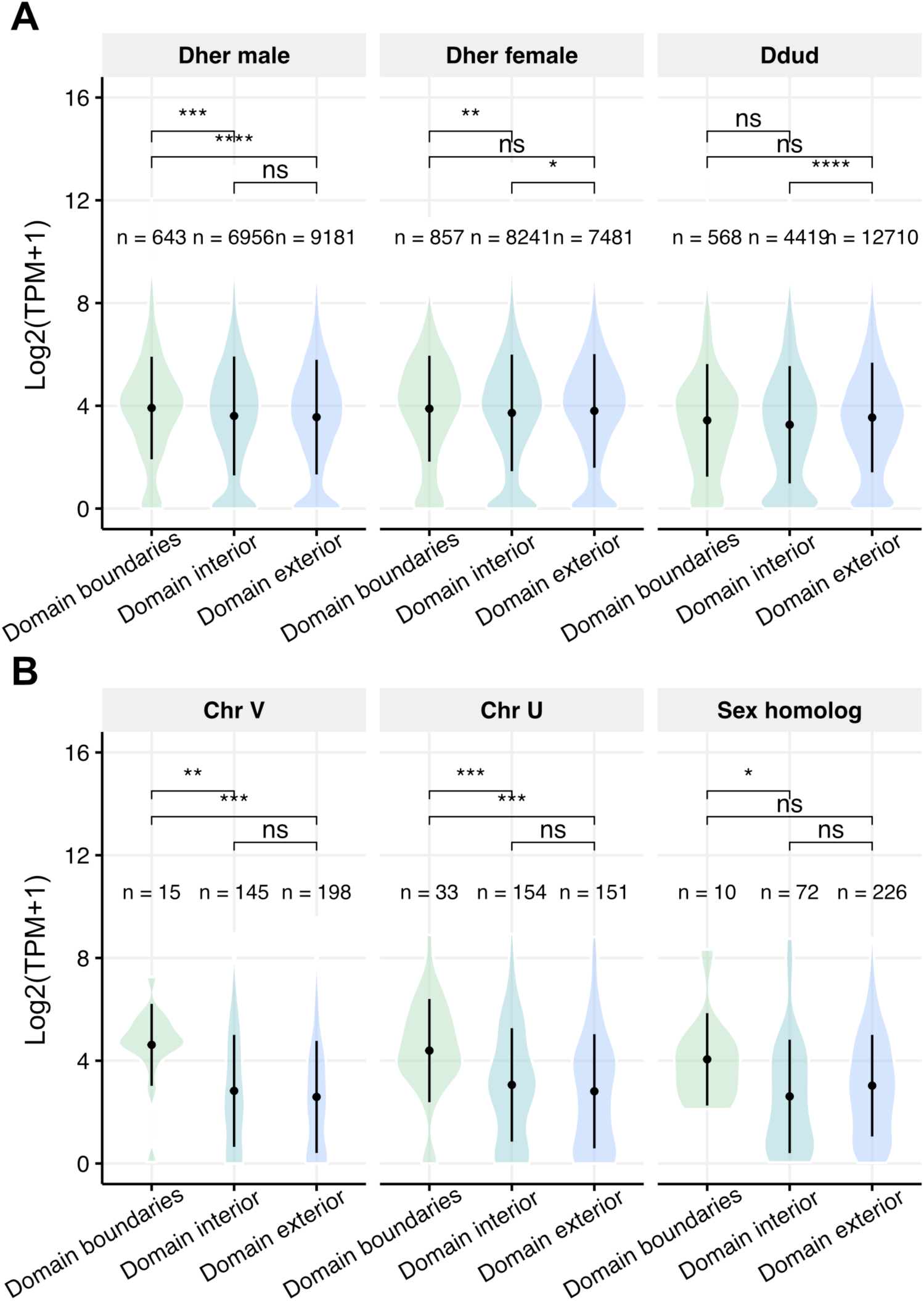
Gene expression profile of chromatin domain for (**A**) whole genome *D. herbacea* (Dher) male female and *D. dudresnayi* (Ddud) and (**B**) sex chromosomes (Chr V and Chr U) and sex-homolog. Numbers above brackets represent the number of genes. Plots show mean ± standard deviation, with violins depicting full data distribution. Significance was determined by a two-sample Wilcoxon rank sum test (****: *p* <= 0.0001, ***: *p* <= 0.001, **: *p* <= 0.01, *: *p* <= 0.05, ns: not significant, *p* > 0.05).

**Fig. S15.**
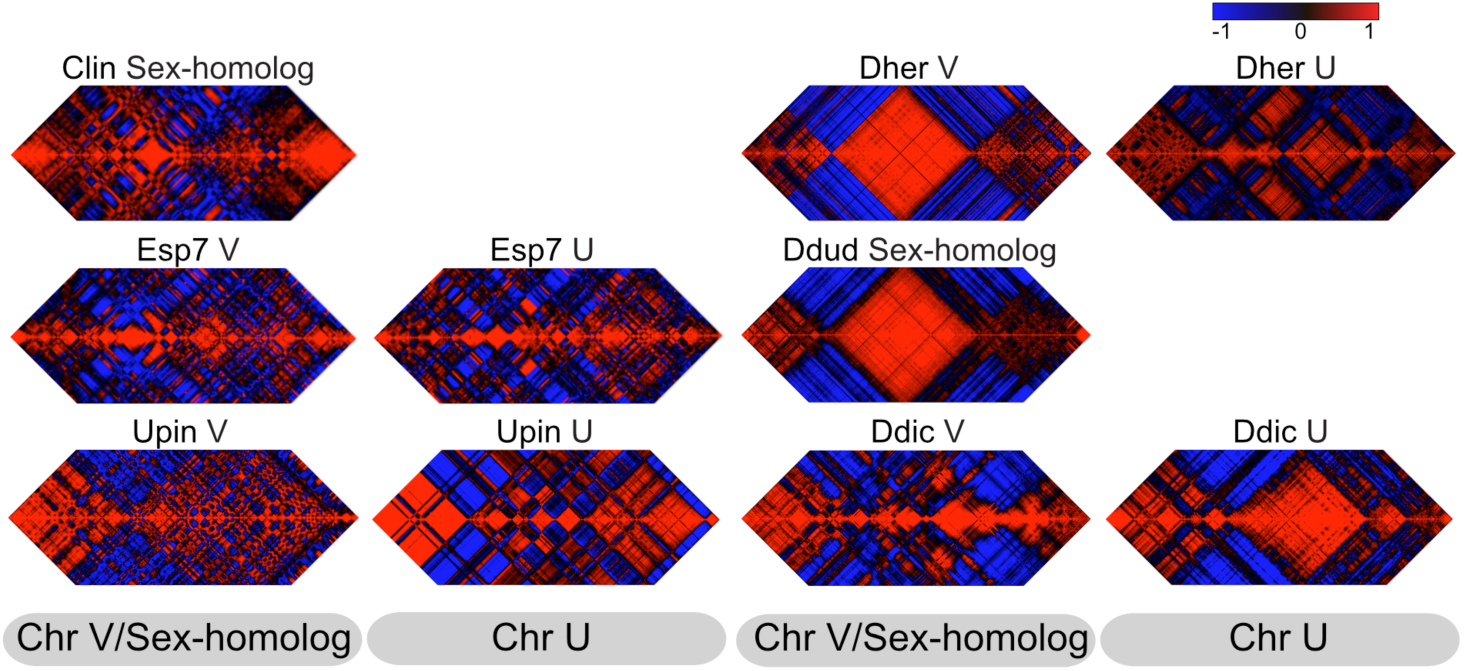
Pearson correlation Hi-C maps at 50k resolution of sex chromosomes and sex-homologs for each species.

**Fig. S16.**
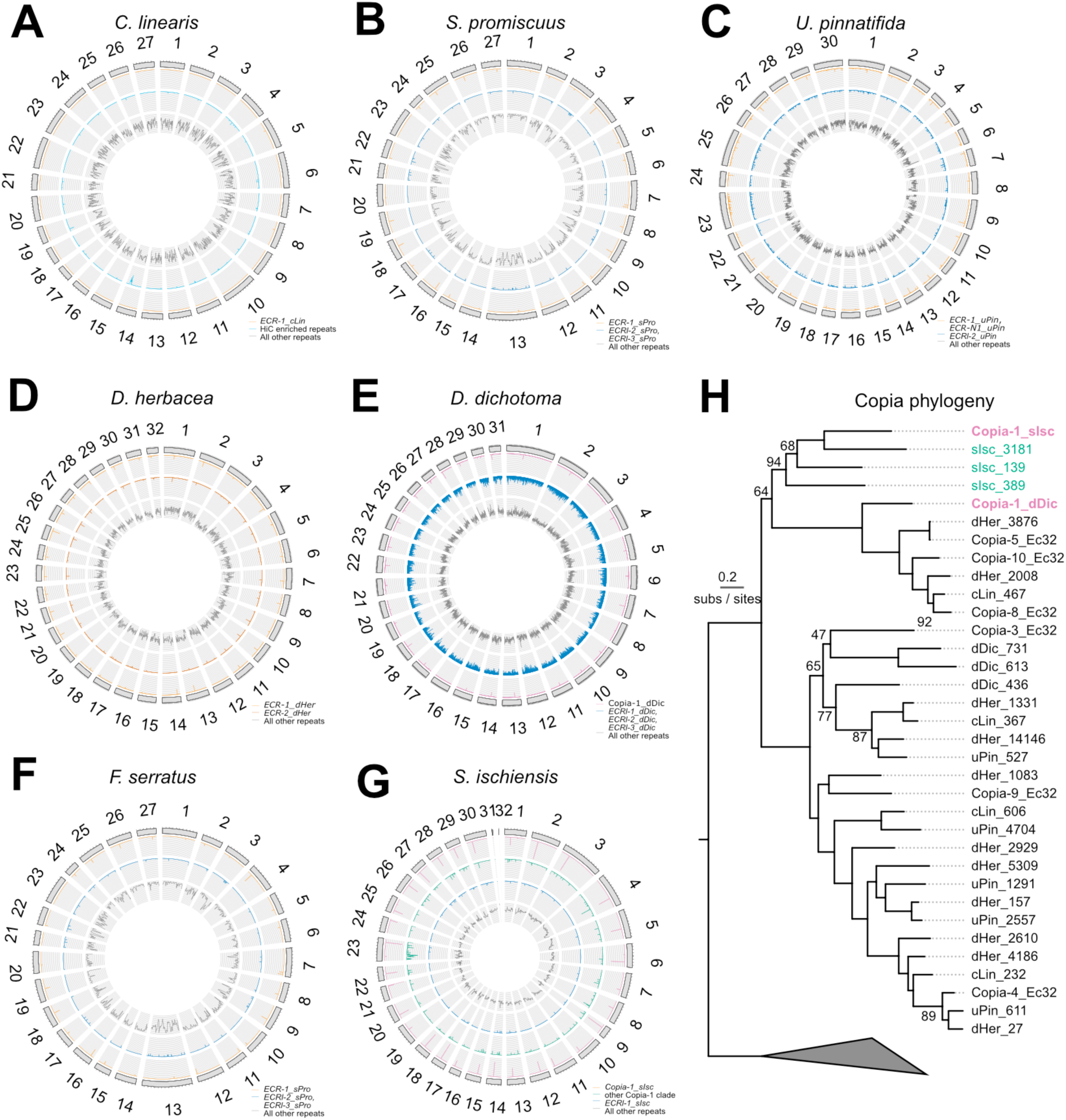
Genome wide distributions of *ECR* and *ECR-like* LTR families and centrophilic Copia families (**A-G**). Genomic proportions for each specific repeat family are in 50 kb windows, whereas all other repeats are in 200 kb windows. Centrophilic retrotransposons have a characteristic single peak per chromosome. (**H**) Maximum likelihood phylogeny of gag and pol protein sequences for Copia families present in the genomes of the species analysed in this study (Ec32 = *Ectocarpus* sp. 7, cLin = *Chordaria linearis*, uPin = *Undaria pinnatifida*, dHer = *Desmarestia herbacea*, dDic = *Dictyota dichotoma*, sIch = *Schizocladia ischiensis*) build under the Q.pfam+R5 model. Centrophilic elements are colored purple and non-centrophilic elements in *S. ischiensis* are colored green (**G**). Ultrafast bootstrap values <95 are shown and the phylogeny is rooted on more distantly related Copia elements from *Ectocarpus* sp. 7.

**Fig. S17.**
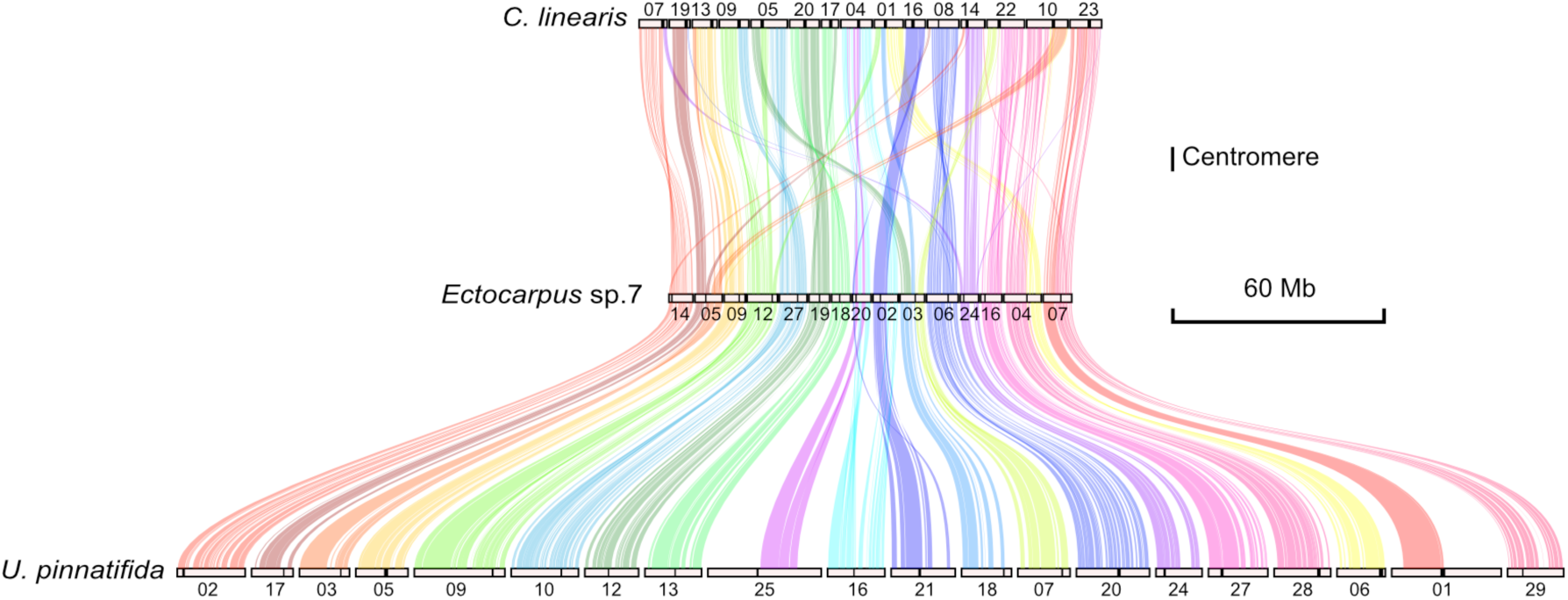
Example shows inter-chromosomal rearrangements events found in *C. linearis*, *Ectocarpus* sp.7 and *U. pinnatifida*, synteny colors were rephased by genome of *U. pinnatifida*, centromeres are plotted as black bars.

**Fig. S18.**
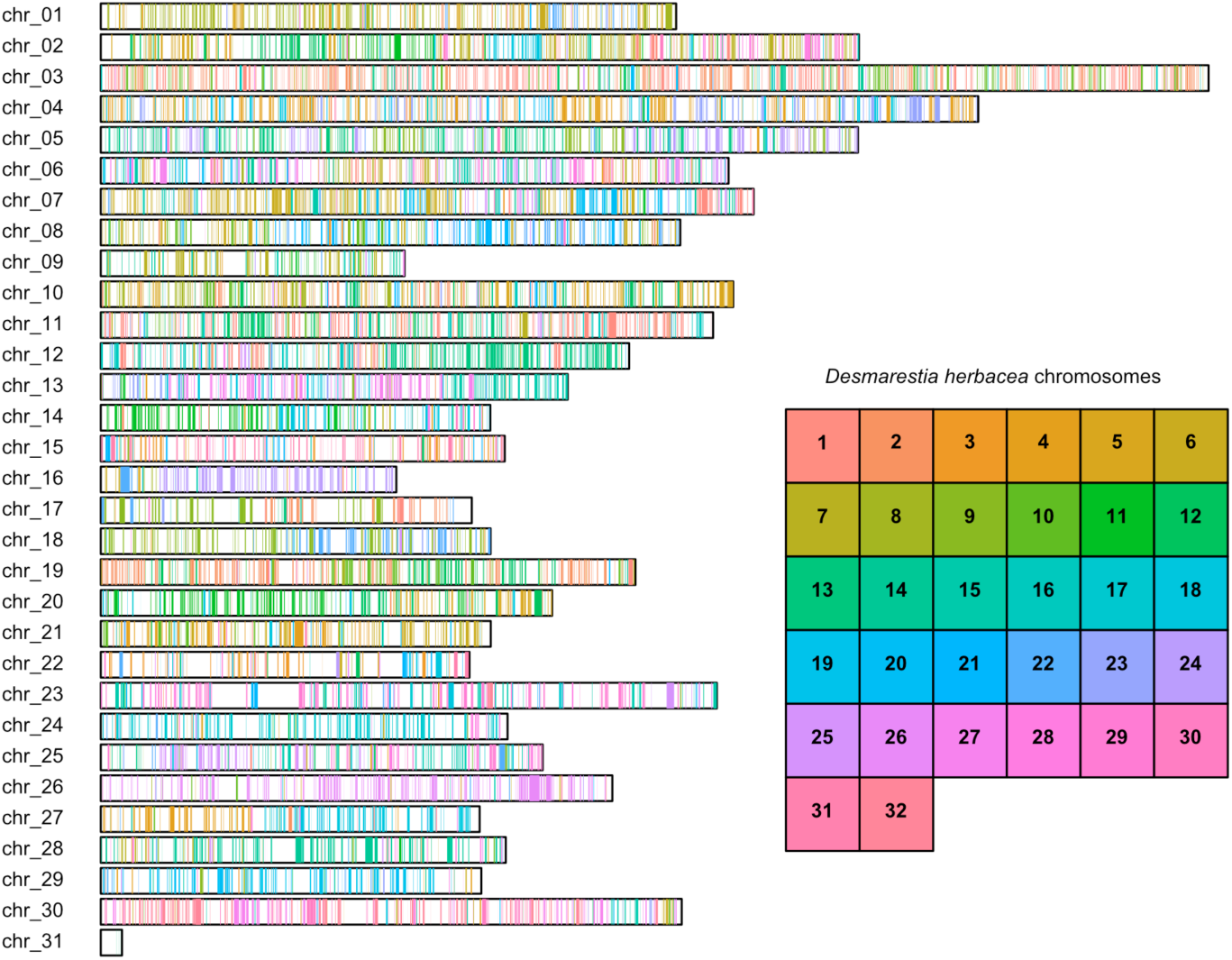
*Schizocladia ischiensis* chromosomes painted by chromosome-of-origin for orthologous genes in *Desmarestia herbacea* (n=8,643).

## Supplemental Tables Legends

**Table S1**. Genomic data used in this study.

**Table S2**. Genome statistics (whole genome assembly + chromosome level assembly + unanchored contigs).

**Table S3**. Gene annotation statistics for species used in this study (male and monoicous data).

**Table S4**. Statistics of number of Hi-C reads for each sample and replicates from Juicer pipeline^101^

**Table S5**: Hi-C resolution used for different analysis across brown algal species

**Table S6**: Pearson correlation of linearly transformed Hi-C matrix at multiple resolutions for orthologous chromosomes between species. Rearranged chromosomes excluded from the analysis.

## Notes

### Competing Interest Statement

The authors have declared no competing interest.

